# Ventral hippocampal projections to the medial prefrontal cortex regulate social memory

**DOI:** 10.1101/461533

**Authors:** Mary L. Phillips, Holly A. Robinson, Lucas Pozzo-Miller

## Abstract

Inputs from the ventral hippocampus (vHIP) to the medial prefrontal cortex (mPFC) have been implicated in several neuropsychiatric disorders. Here, we show that the long-range vHIP-mPFC projection is hyperactive in the *Mecp2* knockout (KO) mouse model of the autism spectrum disorder Rett syndrome, which has deficits in social memory. Chronically mimicking vHIP-mPFC hyperexcitability in wild-type mice impaired social memory, whereas chronic inhibition of mPFC-projecting vHIP neurons in *Mecp2* KO mice rescued social memory deficits; the extent of memory rescue was negatively correlated with the strength of vHIP input to the mPFC. Acute manipulations of the vHIP-mPFC projection also affected social memory in a specific and selective manner, suggesting that proper vHIP-mPFC signaling is necessary to recall social memories. In addition, we identified an altered vHIP-mPFC innervation pattern and increased synaptic strength onto layer 5 pyramidal neurons as contributing factors in aberrant vHIP-mPFC signaling in *Mecp2* KO mice.

## INTRODUCTION

Social interactions are a fundamental part of our daily lives, and impairments in social cognition are key features of multiple neuropsychiatric illnesses. A person or animal must reliably recall previous social interactions to make appropriate social responses and then update the memory with each new encounter. Previous studies have identified a hippocampal network as the neural region that tracks social encounters in both human subjects and mouse models (Hitti and Siegelbaum, 2014; Meira et al., 2018; Okuyama et al., 2016; Tavares et al., 2015). Functional neuroimaging in human subjects has revealed that h i gher covariance between hippocampal activity and changes in social environment reflect better social skills (Tavares et al., 2015). In mouse models, perturbing neuronal activity in both dorsal CA2 and ventral CA1 hippocampal subregions impairs social memory (Hitti and Siegelbaum, 2014; Meira et al., 2018; Okuyama et al., 2016). However, debate remains as to which long-range projections from the hippocampus are required for the formation of social memories.

Autism spectrum disorders (ASDs) are characterized by difficulties in interpreting social situations and a lack of social appropriation (Barendse et al., 2018). A common feature in mouse models of monogenic ASDs is an imbalance in synaptic excitation and inhibition (E/I) in different brain regions (Nelson and Valakh, 2015). Altering the E/I balance in the medial prefrontal cortex (mPFC) in mice mimics autism-like social deficits (Yizhar et al., 2011), and restoring the E/I balance in the *CNTNAP2* knockout (KO) and valproic acid mouse models of ASDs improves their social deficits (Selimbeyoglu et al., 2017; Brumback et al., 2017). Excitatory pyramidal neurons of the ventral hippocampus (vHIP) send long-range projections to the mPFC (Dégenètais et al., 2003; Dembrow et al., 2010; Liu and Carter, 2018; Thierry et al., 2000), and the activity of different populations of mPFC pyramidal neurons are correlated with the novelty of a social target (Liang et al., 2018). Thus, the mPFC is a prime candidate region for the relay of social memory-related signaling from the vHIP. Therefore, we tested whether altering the activity of mPFC-projecting vHIP neurons affects social behavior and memory in a mouse model of the monogenic syndromic ASD Rett syndrome (RTT). We focused on the *Mecp2* KO mouse model of RTT because of the heightened activity in the vHIP (Calfa et al., 2011, 2015) and the hypoactivity of cortical regions (Durand et al., 2012; Kron et al., 2012; Morello et al., 2018; Sceniak et al., 2015; Tomassy et al., 2014), both resulting from opposite changes in the local E/I balance.

Using a combination of unbiased behavioral analyses, selective chemogenetic manipulations with an intersectional genetic approach, high-speed imaging of network activity with voltage-sensitive dyes, intracellular recordings, and dual-color optogenetics, we showed that the long-range vHIP-mPFC projection is hyperactive in RTT mice, which results in social memory deficits. Furthermore, chemogenetic manipulation of mPFC-projecting vHIP neurons in wild-type (WT) and RTT mice correlated with social memory performance in a specific and selective manner. Lastly, these behavioral consequences arose from alterations in the morphology and function of excitatory synapses between vHIP axons and pyramidal neurons in layer 5 of the mPFC.

## RESULTS

### mPFC-projecting vHIP neurons are selectively activated during social encounters

Because both the vHIP and mPFC have been independently implicated in different aspects of social behavior (Hitti and Siegelbaum, 2014; Liang et al., 2018; Meira et al., 2018; Okuyama et al., 2016; Selimbeyoglu et al., 2017; Yizhar et al., 2011), we first tested if the vHIP projection to the mPFC is selectively engaged during social encounters. To identify vHIP neurons based on their long-range projections, we bilaterally injected green RetroBeads into the prelimbic (PL) region of the mPFC and red RetroBeads into the lateral hypothalamus (LH) of male WT and *Mecp2* KO mice at postnatal day 31 (P31), and then we allowed 14 days for RetroBead transfer until P45, when male *Mecp2* KO mice are symptomatic (Chen et al., 2001) (Figures 1A and 1B). We placed mice into a testing chamber, where they were sequentially exposed to either two inanimate objects (two novel fake mice) or to other live mice (a co-housed WT littermate and a novel age-matched WT mouse). Forty-five minutes after the last interaction, we perfused the mice and prepared the brain for quantitative immunohistochemistry of the immediate early gene c-Fos as an estimate of neuronal activity (Figures 1C and 1D). All vHIP neurons in WT mice that interacted with other mice showed higher c-Fos intensities than those in mice that interacted with inanimate objects, irrespective of their projections (p<0.0001, Kruskal-Wallis test with Dunn’s Multiple Comparisons; Figure 1 E). Furthermore, in WT mice that interacted with other mice, mPFC-projecting vHIP neurons had higher c-Fos intensities than those projecting to the LH (n=275 mPFC-projecting neurons in six sections from four mice, n=282 LH-projecting neurons in four sections from three mice; p=0.0098, Kruskal-Wallis test with Dunn’s Multiple Comparisons).

**Figure 1.**
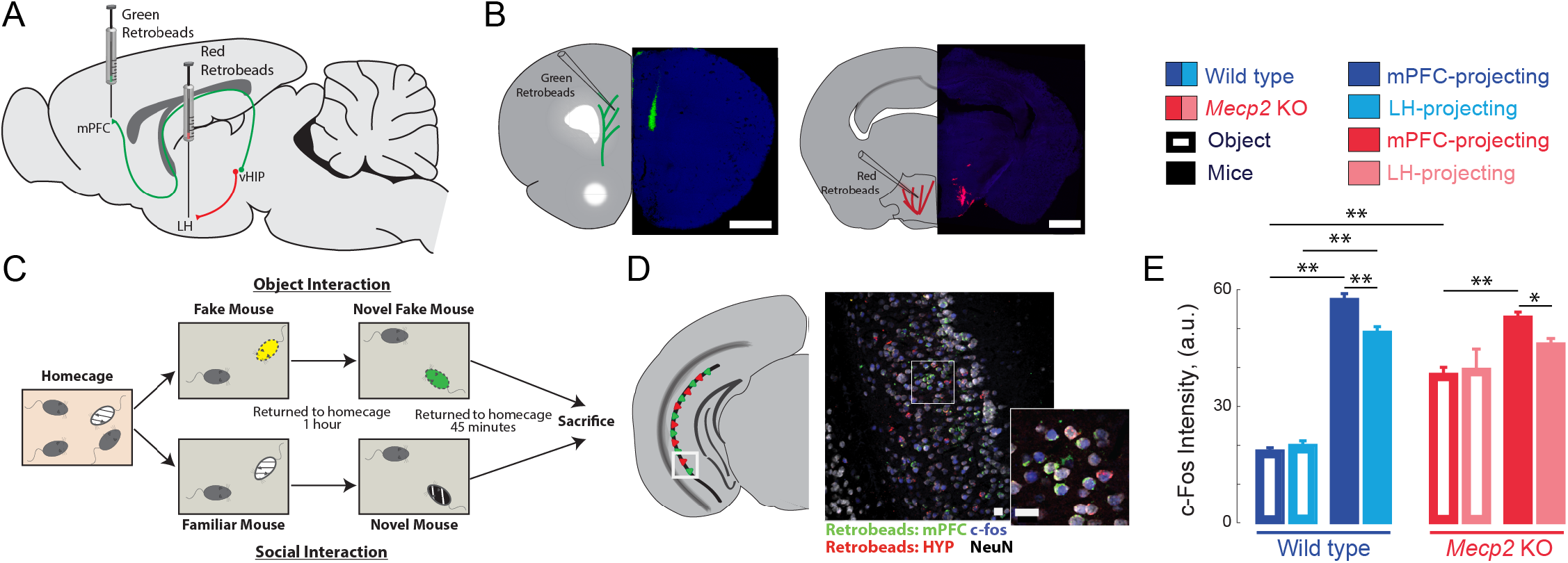
mPFC-projecting vHIP neurons are selectively activated during sequential social encounters with familiar and novel mice. **(A)** Schematic of RetroBead injection for labeling mPFC-or LH-projecting vHIP neurons. **(B)** Injection sites. Scale bar 1 mm. **(C)** Schematic of behavioral paradigm. **(D)** Retrobead labeling and c-Fos immunohistochemistry in vHIP sections. Scale bar 25 μm. **(E)** c-Fos intensity of RetroBead-labeled neurons [n = 163 cells from three sections from two mice (163/3/2) WT mPFC-projecting with object; n = 180/3/2 WT LH-projecting with object; n = 275/6/4 WT mPFC-projecting with mice; n = 282/4/3 WT LH-projecting with mice; n = 105/6/3 *Mecp2* KO mPFC-projecting with object; n = 27/4/3 *Mecp2* KO LH-projecting with object; n = 247/4/3 *Mecp2* KO mPFC-projecting with mice; n = 172/6/3 *Mecp2* KO LH-projecting with mice; WT mPFC object vs. WT LH object, p >0.9999; WT mPFC object vs. WT mPFC mice, p < 0.0001; WT LH object vs. WT LH mice, p < 0.0001; WT mPFC mice vs. WT LH mice, p = 0.0098; KO mPFC object vs. KO LH object, p >0.9999; KO mPFC object vs. KO mPFC mice, p < 0.0001; WT LH object vs. WT LH mice, p > 0.9999; WT mPFC mice vs. WT LH mice, p = 0.0365; WT mPFC object vs. KO mPFC object, p < 0.0001; WT mPFC mice vs. KO mPFC mice, p > 0.9999; Kruskal-Wallis with Dunn’s multiple comparisons test]. Mean ± SEM; * p < 0.05, ** p < 0.01. See also Figure S1.

We obtained similar results in *Mecp2* KO mice, with higher c-Fos intensities in mPFC-projecting vHIP neurons of mice interacting with other mice compared to those interacting with inanimate objects, and higher c-Fos intensities in mPFC-projecting vHIP neurons compared to LH-projecting vHIP neurons in mice interacting with other mice (p<0.05, Kruskal-Wallis test with Dunn’s Multiple Comparisons; Figure 1E). The differences in c-Fos intensities between the object and social conditions were smaller in *Mecp2* KO mice than in WT mice, suggesting a loss of selectivity in vHIP-mPFC signaling likely caused by the higher basal activity in the vHIP of *Mecp2* KO mice (Calfa et al., 2011). Consistent with the lower levels of neuronal activity reported previously (Kron et al., 2012), the PL region of the *Mecp2* KO mPFC had fewer c-Fos-positive neurons than that of WT mice in both the object and social conditions (p<0.05, Kruskal-Wallis test with Dunn’s Multiple Comparisons; Figure S1). However, both WT and *Mecp2* KO mice from the social condition had more c-Fos-positive neurons compared to the object condition, indicating that the mPFC of both genotypes is more robustly activated during a social encounter than by exposure to novel inanimate objects (p<0.001, Kruskal-Wallis test with Dunn’s Multiple Comparisons; Figure S1).

### Atypical social behavior and impaired social memory in *Mecp2* KO mice

To assess social behaviors, we used a three-chamber interaction arena to sequentially test social preference and then social memory. For social preference, we allowed mice to explore either a chamber containing a novel mouse (stranger 1) restrained under an inverted pencil cup or a chamber containing an empty inverted pencil cup (Figure 2A). Both WT and *Mecp2* KO mice spent significantly more time in the chamber containing stranger 1 compared to the empty chamber (n=8 WT mice, p=0.002; n=6 *Mecp2* KO mice, p=0.0106, Student’s paired t-test; Figures 2B and 2C), indicating that social preference is intact in *Mecp2* KO mice. Immediately following the sociability test (within two minutes), we placed a second novel mouse (stranger 2) under the previously empty pencil cup and again allowed mice to explore the chambers. Indicative of social memory for stranger 1 and their preference for novel mice, WT mice spent more time in the chamber with stranger 2 (n=8 WT mice, p=0.0122, Student’s paired t-test; Figures 2B and 2C). However, *Mecp2* KO mice spent similar amounts of time in both chambers (n=6 *Mecp2* KO mice, p=0.768, Student’s paired t-test; Figures 2B and 2C), indicating a deficit in the social memory of the stranger 1 mouse that was encountered two minutes before.

**Figure 2.**
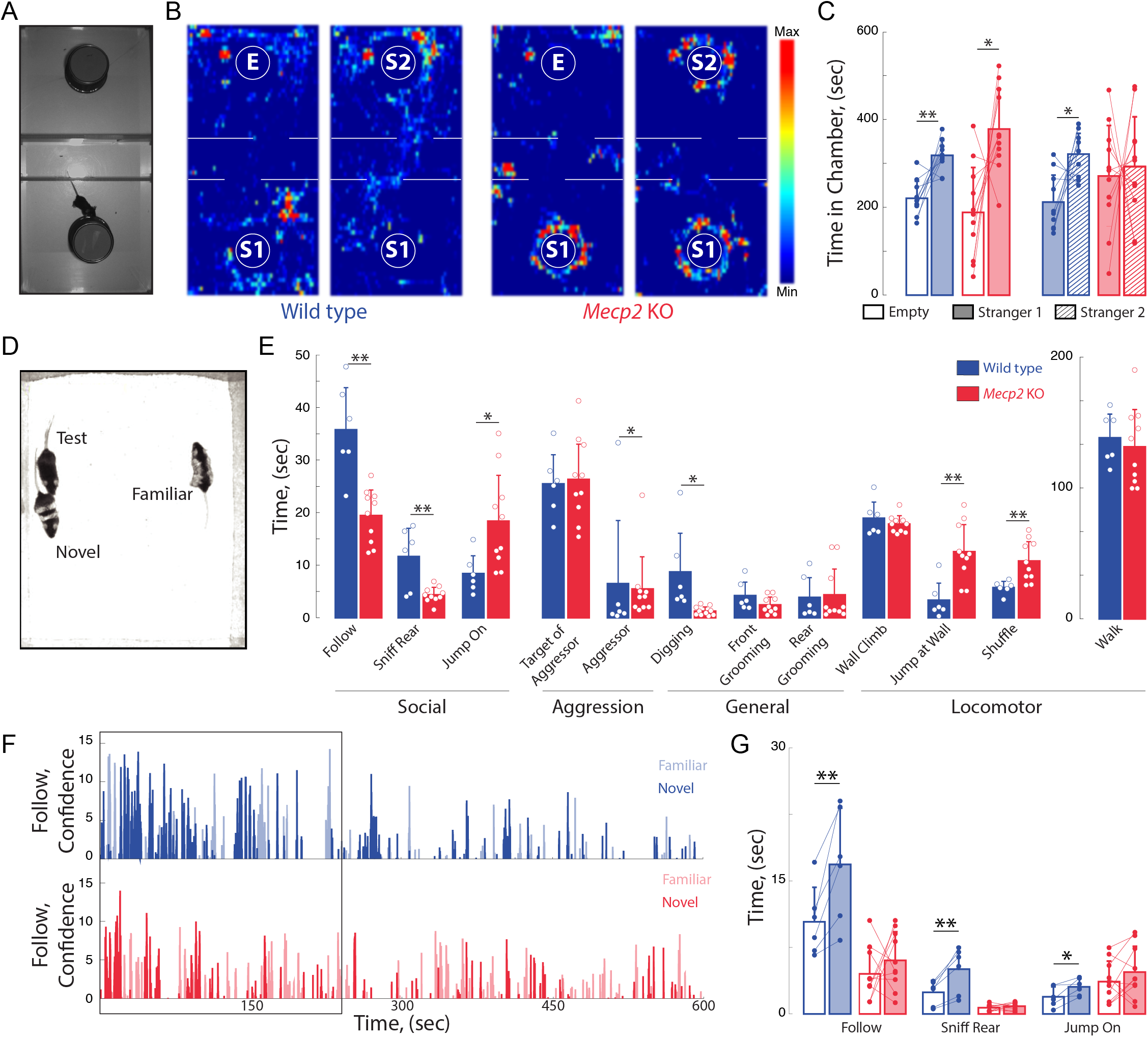
*Mecp2* KO mice have impaired social memory and atypical social interactions. **(A)** Schematic of three-chamber social test. **(B)** Representative heat maps of WT and *Mecp2* KO mice during the three-chamber social test. **(C)** Time spent in chambers with the empty inverted pencil cup or cup containing stranger 1 or stranger 2 (n = 10 WT mice; n = 11 *Mecp2* KO mice; WT empty vs. WT social, p = 0.0020; *Mecp2* KO empty vs. *Mecp2* KO social, p = 0.0106; WT familiar vs. WT novel, p = 0.0122; *Mecp2* KO familiar vs. *Mecp2* KO novel, p = 0.7680; Student’s paired t-test). **(D)** Image of unrestricted social memory paradigm. **(E)** Time spent performing different behaviors using automated scoring of unrestricted social encounters (n = 6 WT mice; n = 6 *Mecp2* KO mice; Follow, p = 0.0003; Rear Sniff, p = 0.0020; Jump On, p = 0.0250; Target of Aggressor, p = 0.5696; Aggressor, p = 0.0378; Digging, p = 0.0101; Grooming Front, p = 0.1269; Grooming Rear, p = 0.8426; Wall Climb, p = 0.3844; Wall Jump, p = 0.0025; Shuffle, p = 0.0087; Walk, p = 0.6268; Student’s t-test for all except Aggressor, which used Kolmogorov-Smirnov test). **(F)** Computer confidence scores in following behavior over the course of representative videos separated by target: novel mouse in bold and familiar opaque, first four minutes boxed. **(G)** Time the test mouse spent following, sniffing rear, or “jumping on” either the novel or familiar mouse during the first four minutes of the video (n = 6 WT; n = 6 KO mice; WT Follow, p = 0.0098; WT Rear Sniff, p = 0.0056; WT Jump On, p = 0.0289; *Mecp2* KO Follow p = 0.4316, *Mecp2* KO Rear Sniff, p = 0.3281; *Mecp2* KO Jump On, p = 0.2349; Student’s paired t-test except for *Mecp2* KO Follow, which used Wilcoxon paired test). Mean ± SD; * p < 0.05, ** p < 0.01.

To avoid potential confounds arising from testing social interactions with restrained mice, we implemented a behavioral assay in which WT or *Mecp2* KO mice freely interacted simultaneously for 10 minutes with both a co-housed WT littermate and a novel age-matched WT mouse (Figure 2D). Unbiased scoring of social interactions by the machine-learning based Janelia Automatic Animal Behavior Annotator (*JAABA*) (Kabra et al., 2013; Ohayon et al., 2013; Robie et al., 2017) revealed that WT mice mainly engaged in following behavior. By contrast, *Mecp2* KO mice followed and sniffed other mice less than WT mice, and engaged in an atypical “jumping on” behavior (Figure 2E). In addition, *Mecp2* KO mice displayed less digging, more wall jumping, and more shuffled walking (n=6 WT mice, n=10 *Mecp2* KO mice, p<0.05, Student’s t-test; Figure 2E). As a critical control for social interactions in the unrestricted arena and the three-chamber test, WT and *Mecp2* KO mice spent a comparable amount of time walking (p=0.6268, Student’s t-test; Figure 2E). Overall, these data indicate that, while *Mecp2* KO mice do display interest in other mice, they do so in an atypical manner.

During the first four minutes of the assay, WT mice preferentially interacted with the novel mouse across all measured social behaviors, including following, sniffing, and “jumping on”, after which social interaction declined regardless of target (n=6 mice, p<0.05, Student’s paired t-test Figure 2F). However, *Mecp2* KO mice showed no significant preference between the familiar and novel mouse (n=10 mice, p>0.05, Student’s paired t-test; Figure 2G), consistent with previously observed deficits in social memory (see Figure 2C).

### Increased influence of vHIP input on the mPFC network in *Mecp2* KO mice

Because mPFC-projecting vHIP neurons are activated during social encounters and *Mecp2* KO mice have impaired social memory and atypical social behaviors, we characterized vHIP inputs to the mPFC in *Mecp2* KO mice at the functional level. To identify hippocampal fibers in *ex vivo* slices of the mPFC, we injected the fluorescent tracer dextran-Alexa-594 into the vHIP and cut brain slices at a 10° angle from the coronal plane (Parent et al., 2010) (Figure 3A). We evoked single field excitatory postsynaptic potentials (fEPSPs) with a theta-glass electrode placed in the fluorescently labeled vHIP fiber bundle and imaged voltage-sensitive dye (VSD) signals, which are directly proportional to the amplitude of fEPSPs and follow their kinetics (Grinvald et al., 1988) (Figures 3B and 3C). The amplitudes of VSD signals evoked by stimulation of vHIP afferents were larger in mPFC slices from *Mecp2* KO mice compared to WT littermates at a range of stimulation intensities [n=11 slices from 7 WT mice, n=11/5 *Mecp2* KO mice, p=0.0470, Two-way analysis of variance (ANOVA); Figures 3D and 3E]. The spatiotemporal spread of VSD signals throughout the mPFC slice was similar overall between *Mecp2* KO and WT mice (p=0.4529, Two-way ANOVA; Figure 3F), although the spatial spread was significantly larger at lower stimulation intensities in slices from *Mecp2* KO mice (p=0.0133, Mann-Whitney test; Figure 3G). Stimulating intracortical fibers with another theta glass electrode placed in layer 2/3 of the same cortical column evoked VSD amplitudes of comparable amplitude in slices from *Mecp2* KO and WT mice (n=11 slices from 7 WT mice, n=11/5 *Mecp2* KO mice, p=0.0540; Two-way ANOVA; Figures 3H and 3I). By contrast, the spatiotemporal spread of VSD signals evoked by intracortical stimulation was significantly smaller in slices from *Mecp2* KO mice (p=0.0498, Two-way ANOVA; Figures 3J and 3K). The amplitude and spatial spread of VSD signals evoked by vHIP stimulation were 72% and 71%, respectively, of those evoked by intracortical stimulation in mPFC slices from WT mice. However, the amplitude and spatial spread of VSD signals evoked by vHIP stimulation were 95% and 96% of those evoked by intracortical stimulation in *Mecp2* KO slices, which reflects both larger vHIP-evoked signals and smaller responses to intracortical stimulation (Figures 3L-3N). These data indicate that vHIP fibers drive hyperactivation of the mPFC network in *Mecp2* KO mice, in contrast to the hypoactivation driven by intracortical stimulation, suggesting that vHIP inputs are overrepresented in the mPFC network of *Mecp2* KO mice.

**Figure 3.**
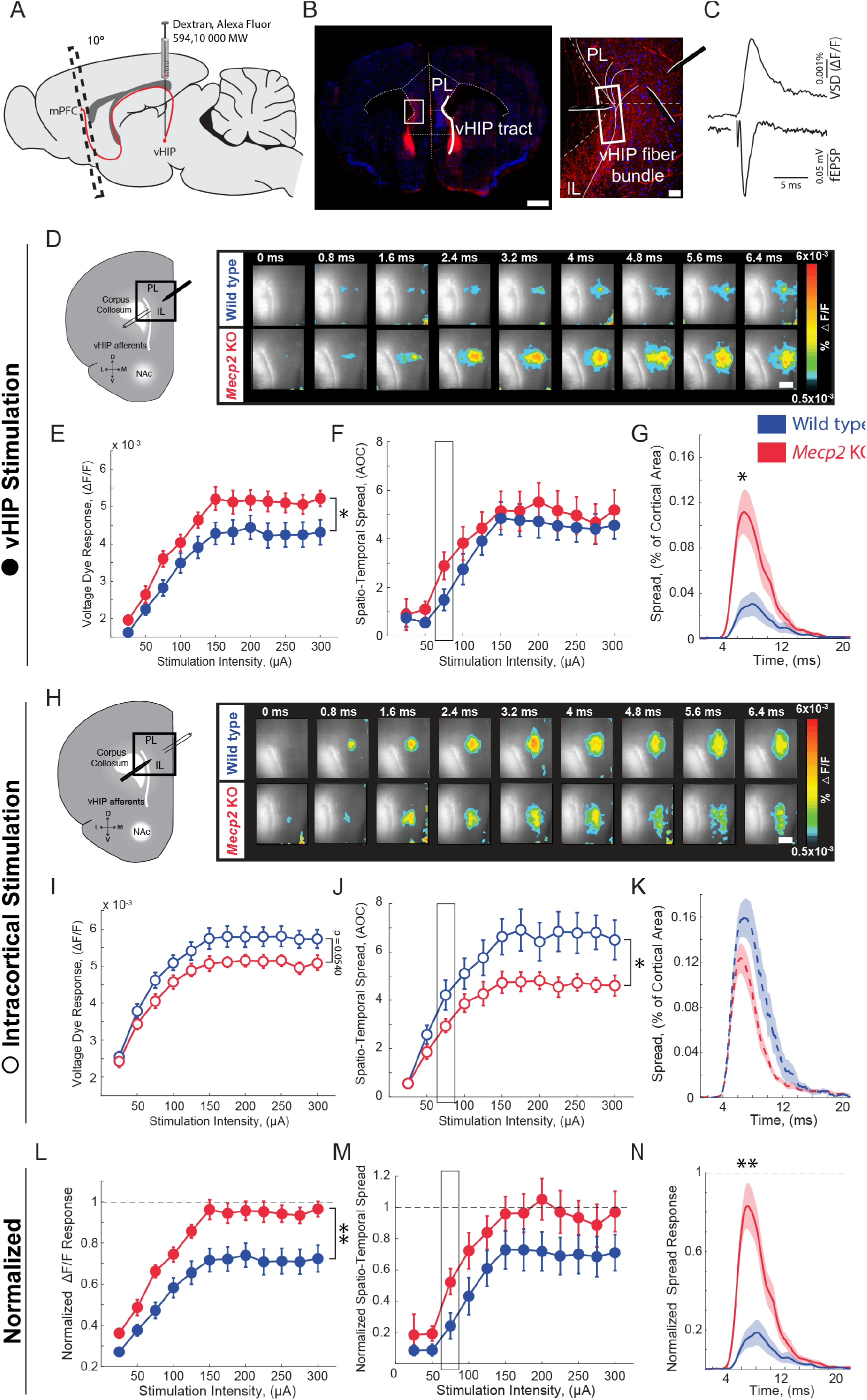
Increased influence of vHIP axons in the mPFC of *Mecp2* KO mice. **(A)** Schematic of dextran injection into the vHIP. **(B)** Visualization of vHIP fibers in mPFC slices. Scale bar 0.5 mm; inset 100 μm. **(C)** VSD responses are proportional to the amplitude and follow the kinetics of fEPSPs. **(D)** Representative VSD responses evoked by stimulation of fluorescently-labelled vHIP fibers. Scale bar 100 μm (**D & H**). **(E-G)** Input-output relationship of peak VSD responses (**E**, p = 0.047, Two-way ANOVA), spatio-temporal spread (**F**, p = 0.4529, Two-way ANOVA), and spread over time (**G**, p = 0.0133, Mann-Whitney) (at 75 μA intensity) evoked by vHIP fiber stimulation. **(H)** Representative VSD responses evoked by intracortical stimulation. **(I-K)** Input-output relationship of peak VSD responses (**I**, p = 0.4553, Two-way ANOVA), spatio-temporal spread (**J**, p = 0.0498, Two-way ANOVA), and spread over time (**K**, p = 0.1025, Student’s t-test) (at 75 μA intensity) evoked by intracortical stimulation. **(L-N)** Peak VSD responses (**L**, p = 0.0015, Two-way ANOVA), spatiotemporal spread (**M**, p = 0.0767, Two-way ANOVA), and spread over time (**N**, p = 0.0002, Mann-Whitney) evoked by vHIP fiber stimulation normalized to those evoked by intracortical stimulation. [n = 11 slices from 7 mice (11/7) WT mice; n = 11/5 *Mecp2* KO]. Spatio-temporal spread = AOC created by spread of the cortical area (% of total) and time (ms).

Considering the role of long-term synaptic plasticity in memory, we tested the ability of excitatory vHIP-mPFC synapses to undergo long-term potentiation (LTP) in slices from *Mecp2* KO mice, because previous studies have described plasticity at these synapses in rats and mice *in vivo* (Izaki et al., 2003, 2001; Laroche et al., 2000). In mPFC slices from WT mice, high-frequency stimulation of vHIP afferents evoked a significant potentiation of the spatiotemporal spread of VSD signals, which persisted up to 45 min (n=10 slices from 7 mice, p=0.0013, Student’s paired t-test) and was sensitive to the N-methyl-D-aspartate (NMDA) receptor antagonist APV (100 μM) (n=4 slices from 4 mice, p=0.9205; Student’s paired t-test; Figure S2). However, mPFC slices from *Mecp2* KO mice showed only a short-term enhancement of the spatiotemporal spread of VSD signals, which quickly decayed back to baseline levels (n=9 slices from 5 mice, p=0.2705; Student’s paired t-test; Figure S2). These data demonstrate an impairment of LTP at excitatory vHIP-mPFC synapses, similar to that previously reported at CA3-CA1 synapses in hippocampal slices of *Mecp2* KO mice (Li et al., 2016).

### Selective chemogenetic manipulation of mPFC-projecting vHIP neurons regulates social memory

To causally link the enhanced vHIP input to the mPFC with the deficits in social behavior in *Mecp2* KO mice, we used an intersectional genetic approach to express “designer receptors exclusively activated by designer drugs” (DREADDs) selectively in mPFC-projecting vHIP neurons, and then modulate their activity with the designer ligand clozapine-N-oxide (CNO) (Armbruster et al., 2007; Boender et al., 2014). Briefly, we injected a retrogradely transported canine adenovirus-2 (CAV-2) expressing Cre recombinase (Cre; CAV-2-Cre) bilaterally into the mPFC. We then injected adeno-associated virus serotype 8 (AAV8) expressing either excitatory (hM3Dq) or inhibitory (hM4Di) DREADDs from a Cre-dependent double-floxed inverse open reading frame (DIO) (Hnasko et al., 2006; Kremer et al., 2000) bilaterally in the vHIP of WT and *Mecp2* KO mice at P20 (Figure 4A). Control mice injected with CAV2-Cre and AAV8-DIO-mCherry were also treated with CNO to account for potential peripheral conversion of CNO into clozapine (Gomez et al., 2017). This intersectional approach resulted in sparse labeling of pyramidal neurons in the ventral CA1 region with their axons projecting to the mPFC (Figure 4B). To chronically modulate the vHIP-mPFC circuit, we delivered CNO via the drinking water (5 mg/kg/day) (Carvalho Poyraz et al., 2016) beginning at P34 and continuing until we used mice for experiments (Figure 4A). At the age of viral injections and the start of CNO treatment, *Mecp2* KO mice lack the behavioral and cellular features that will develop into Rett-like symptoms after P45 (Calfa et al., 2011; Durand et al., 2012; Tomassy et al., 2014). In addition, we confirmed that P20-25 *Mecp2* KO mice performed at WT levels in terms of social memory (Figure S3), VSD signals in mPFC evoked by either vHIP or intracortical stimulation (Figure S4), and LTP at vHIP-mPFC synapses (Figure S4). Notably, the amplitude of vHIP-evoked VSD signals in mPFC slices from *Mecp2* KO mice did not show the typical developmental reduction between P20-25 and P45-50 observed in WT slices, resulting in significantly larger responses in symptomatic *Mecp2* KO mice compared to age-matched WT mice (Figure S5), similar to CA3-evoked VSD responses in CA1 of hippocampal slices (Calfa et al., 2011).

**Figure 4.**
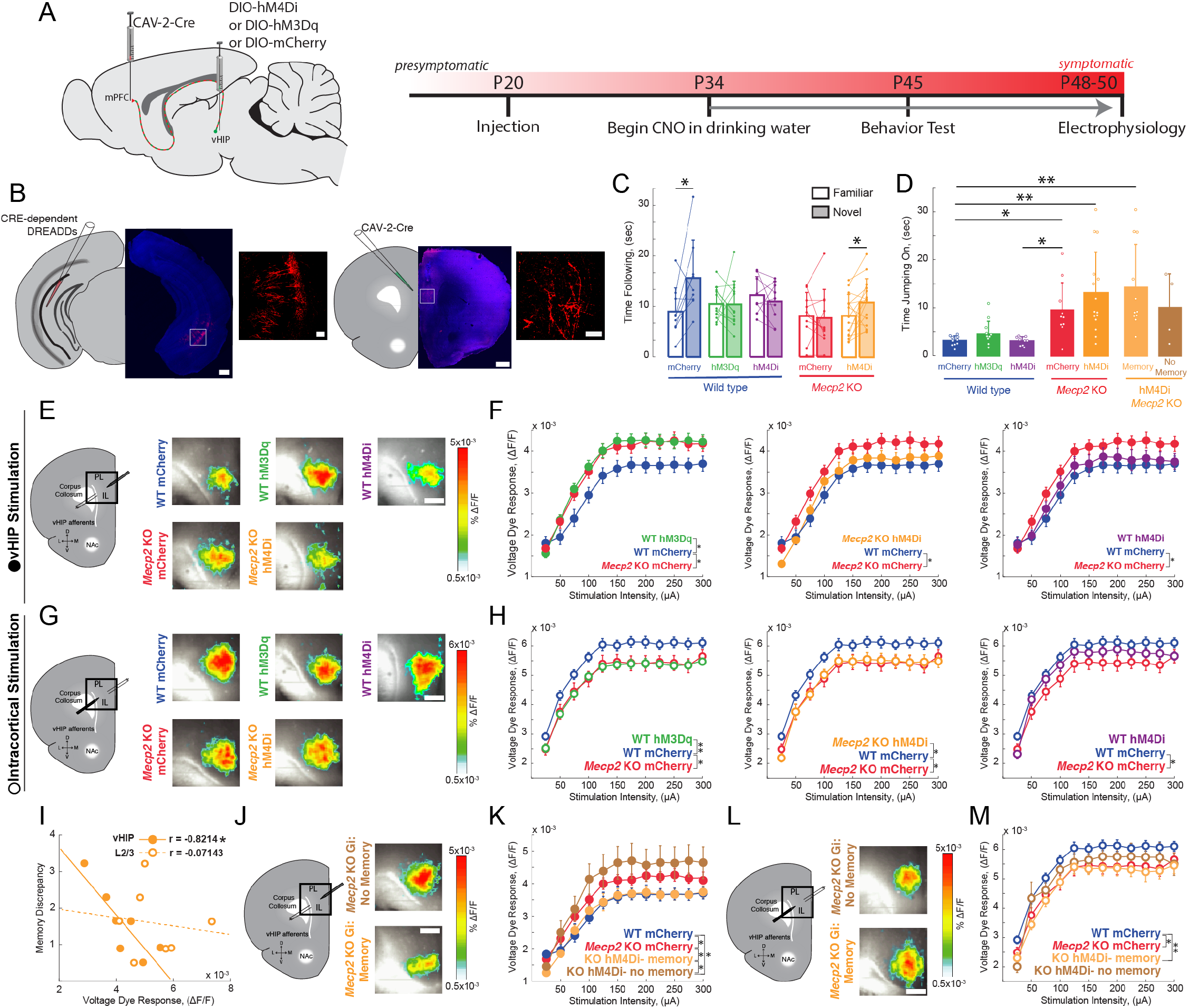
Activity of mPFC-projecting vHIP neurons modulates social memory in WT and *Mecp2* KO mice. **(A)** Schematic of CAV2-Cre and DREADD injections and experimental timeline. **(B)** Injection sites show sparse mCherry labeling of vHIP neurons with identifiable axons in the mPFC. Scale bar 500 μm large, 100 μm inset. **(C)** Time spent following either familiar or novel mice in unrestricted social interaction, scored by *JAABA* (n = 9 mCherry WT mice, p = 0.0488, Wilcoxon paired test; n = 12 hM3Dq WT mice, p = 0.9647, Student’s paired t-test; n = 9 hM4D1 WT mice, p = 0.401, Student’s paired t-test; n = 10 mCherry *Mecp2* KO, p = 0.7871, paired Student’s t-test; n = 15 *Mecp2* KO mice, p = 0.026, Student’s paired t-test). **(D)** Time spent “jumping on” other mice during unrestricted social interaction (n = 9 mCherry WT mice; n = 12 hM3Dq WT mice; n = 9 hM4D1 WT mice; n = 10 mCherry *Mecp2* KO; n = 15 hM4D1 *Mecp2* KO mice; mCherry WT vs. mCherry *Mecp2* KO, p = 0.0215; hM3Dq WT vs. mCherry WT, p > 0.9999; hM3Dq WT vs. hM4D1 WT, p > 0.9999; hM3Dq WT vs. mCherry *Mecp2* KO, p = 0.2119; hM4D1 WT vs. mCherry WT, p > 0.9999; hM4D1 WT vs. mCherry *Mecp2* KO, p = 0.0194; mCherry *Mecp2* KO vs. hM4D1 (All) *Mecp2* KO, p > 0.9999; hM4D1 (All) *Mecp2* KO vs. mCherry WT, p = 0.0016; Memory hM4D1 vs. mCherry KO, p > 0.9999; Memory hM4D1 vs. mCherry WT, p = 0.0012; No Memory hM4D1 vs. mCherry *Mecp2* KO, p > 0.9999; No Memory hM4D1 vs. mCherry WT, p = 0.8023; Memory hM4D1 vs. No Memory hM4D1, p > 0.9999, Kruskal-Wallis test with Dunn’s multiple corrections). **(E)** Representative VSD responses evoked by vHIP fiber stimulation in CNO-treated mice. Scale bar 200 μm. **(F)** Input-output relationships of peak VSD responses evoked by vHIP fiber stimulation. (mCherry WT vs. mCherry *Mecp2* KO, p = 0.0456; hM3Dq WT vs. mCherry WT, p = 0.0254; hM3Dq WT vs. mCherry *Mecp2* KO, p = 0.8351; mCherry WT vs. hM4D1, p = 0.5715; hM4D1 WT vs. mCherry *Mecp2* KO, p = 0.1771; hM4D1 *Mecp2* KO vs. mCherry WT, p = 0.677l; hM4D1 *Mecp2* KO vs. mCherry *Mecp2* KO, p = 0.1095; Two-way ANOVA). **(G)** Representative VSD responses evoked by intracortical stimulation in CNO-treated mice. **(H)** Input-output relationships of peak VSD responses evoked by intracortical stimulation (mCherry WT vs. mCherry *Mecp2* KO, p = 0.0327; hM3Dq WT vs. mCherry WT, p = 0.0023; hM3Dq WT vs. mCherry *Mecp2* KO, p = 0.9193; mCherry WT vs. hM4D1, p = 0.2099; hM4D1 WT vs. mCherry *Mecp2* KO, p = 0.2417; hM4D1 *Mecp2* KO vs. mCherry WT, p = 0.0313; hM4D1 *Mecp2* KO vs. mCherry *Mecp2* KO, p = 0.9045; Two-way ANOVA). **(I)** Correlation between social memory indices to VSD responses evoked by either vHIP fiber (closed circles) or intracortical stimulation (open circles) (n = 7 slices from 7 hM4D1 *Mecp2* KO mice; Spearman r correlation; r = −0.8214, p = 0.0341 vHIP fiber; r = −0.07143, p = 0.9063 intracortical). **(J-K)** VSD input-output relationships evoked by vHIP fiber stimulation in slices from hM4D1 *Mecp2* KO mice with intact or impaired social memory (Memory vs. mCherry WT, p = 0.6377; Memory vs. mCherry *Mecp2* KO, p = 0.0427; No Memory vs. mCherry WT, p = 0.0115; No Memory vs. mCherry *Mecp2* KO, p = 0.1297; Memory vs. No Memory, p = 0.0183; Two-way ANOVA). **(L-M)** VSD input-output relationships evoked by intracortical stimulation in slices from hM4D1 *Mecp2* KO mice with intact or impaired social memory (Memory vs. mCherry WT, p = 0.009; Memory vs. mCherry *Mecp2* KO, p = 0.07679; No Memory vs. mCherry WT, p = 0.291, No Memory vs. mCherry *Mecp2* KO, p = 0.6723; Memory vs. No Memory, p = 0.4187; Two-way ANOVA). **(E-M)** n = 18 slices from 9 mCherry (18/9) WT mice; n = 12/8 hM3Dq WT mice; n = 16/9 hM4D1 WT mice; n = 17/8 mCherry *Mecp2* KO mice; n = 20/10 hM4D1 *Mecp2* KO mice; n = 11/6 hM4D1 *Mecp2* KO memory mice; n = 6/4 hM4D1 *Mecp2* KO no memory mice). **(C-D**) Mean ± SD**; (F,H,K,M)** Mean ± SEM; * p < 0.05, ** p < 0.01. See also Figure S6.

When tested in the unrestricted social assay, P45 WT mice expressing only the marker mCherry in mPFC-projecting vHIP neurons and treated with CNO for 11 days, followed the novel mouse more than the familiar littermate (n=9, p=0.0488; Wilcoxon paired test; Figure 4C), similar to nonsurgical, untreated WT mice. By contrast, CNO-treated WT mice expressing the excitatory DREADD hM3Dq in mPFC-projecting vHIP neurons (to mimic vHIP hyperactivity in *Mecp2* KO mice) did not discriminate between the novel mouse and the familiar littermate (n=12, p=0.9647, Student’s paired t-test; Figure 4C), indicating a deficit in social memory resembling that of *Mecp2* KO mice. Chronic inhibition of mPFC-projecting vHIP neurons with the inhibitory DREADD hM4Di also impaired social memory in WT mice (n=9, p=0.4010, Student’s paired t-test) (Figure 4C), underscoring the role of this long-range projection in social memory.

To further define the consequences of altered vHIP-mPFC signaling in social behaviors, we chronically inhibited mPFC-projecting vHIP neurons in *Mecp2* KO mice with the inhibitory DREADD hM4Di and CNO administration from P34 until P45 to selectively reduce hyperactivation of this circuit. This manipulation was sufficient to increase the time following the novel mouse compared to the familiar littermate in 73% of the treated *Mecp2* KO mice, resulting in a significant preference for the novel mouse and indicating a rescue of social memory (n=15, p=0.0260, Student’s paired t-test). By contrast, control CNO-treated *Mecp2* KO mice expressing mCherry did not discriminate between the novel mouse and the familiar littermate (n=10, p=0.7871, Student’s paired t-test; Figure 4C), similar to nonsurgical, untreated *Mecp2* KO mice. In addition, there were no changes in the types of social behaviors displayed by CNO-treated mice expressing DREADDs in mPFC-projecting vHIP neurons. For example, the amount of time performing the atypical “jumping on” behavior did not differ between hM4Di-expressing and mCherry control *Mecp2* KO mice (p>0.9999 within genotypes; p<0.05 between genotypes; Kruskal-Wallis test with Dunn’s Multiple Comparisons; Figure 4D), suggesting that the vHIP-mPFC projection plays a specific role in the memory aspect of social interactions. Other than a small reduction of walking time in hM3Dq-expressing WT mice (p=0.0296, One-way ANOVA with Tukey’s Multiple Comparisons), there were no differences in grooming behavior, locomotion, or anxiety-like behaviors between DREADD-expressing WT and *Mecp2* KO mice and their mCherry-expressing controls after CNO treatment (p>0.05; Figure S6).

After allowing 3-4 days for the potential effects of behavioral testing to fade, we prepared *ex vivo* mPFC slices from CNO-treated DREADD-and mCherry-expressing mice for VSD imaging. Consistent with their deficit in social memory, WT mice expressing the excitatory DREADD hM3Dq in mPFC-projecting vHIP neurons had larger vHIP-induced VSD signals in mPFC slices compared to mCherry-expressing WT controls (n=18 slices from 9 mCherry WT mice; n=12/8 hM3Dq WT; n=17/8 *Mecp2* KO mice; p=0.0254; p=0.8351; Two-way ANOVA; Figures 4E and 4F), resembling those observed in mCherry-expressing *Mecp2* KO controls. Surprisingly, inhibiting mPFC-projecting vHIP neurons with hM4Di did not affect vHIP-induced VSD signals in mPFC slices of WT mice (Figures 4E and 4F). While unexpected, we cannot rule out the possibility that other afferents may have increased their input to the mPFC, as reported previously (Guirado et al., 2015), or that other homeostatic mechanisms might have maintained proper activity levels in the mPFC. Interestingly, we still observed impaired memory performance, suggesting that dysfunction of the vHIP-mFPC pathway is sufficient to impair memory formation in the absence of large-scale changes to the mPFC network. As a control, mCherry expression in mPFC-projecting vHIP neurons followed by CNO treatment did not alter the difference in the amplitude of vHIP-induced VSD signals in mPFC slices between WT and *Mecp2* KO mice (Figure 4F), which resemble those in nonsurgical, untreated mice (see Figure 3E).

Because 27% of *Mecp2* KO mice expressing the inhibitory DREADD hM4Di in mPFC-projecting vHIP neurons did not show an improvement of social memory in the unrestricted test (see Figure 4C), we correlated their social discrepancy scores (higher scores indicate more time spent following the novel mouse vs. the familiar one) with the amplitude of vHIP-induced VSD signals in mPFC slices; this analysis uncovered a statistically significant negative correlation (n=15, r= −0.8214 p=0.0341, Spearman r correlation; Figure 4I). Furthermore, vHIP-induced VSD signals in mPFC slices from *Mecp2* KO mice that showed improved social memory after expression of hM4Di in mPFC-projecting vHIP neurons were smaller than those in mCherry-expressing *Mecp2* KO controls (n=11/6, p=0.0427, Two-way ANOVA; Figures 4J and 4K), resembling those in mCherry-expressing WT mice. By contrast, vHIP-induced VSD signals in mPFC slices from *Mecp2* KO mice that still showed deficits in social memory after expression of hM4Di in mPFC-projecting vHIP neurons were comparable to those in mCherry-expressing *Mecp2* KO controls, and significantly larger than those in mCherry-expressing WT mice (p=0.1297; p=0.0115; Two-way ANOVA; Figures 4J and 4K). Other than social memory discrepancy, we observed no behavioral differences between the improved and impaired memory hM4Di *Mecp2* KO groups (Figures 4C, 4D, and S6).

Selective chemogenetic excitation of mPFC-projecting vHIP neurons in WT mice from P34 to P45 also affected VSD signals evoked by intracortical stimulation in layer 2/3 of mPFC slices. VSD responses in hM3Dq-expressing WT mice were significantly smaller than those in mCherry-expressing WT mice, and resemble those in mCherry-expressing *Mecp2* KO mice (n=18 slices from 9 mCherry WT mice; n=12/8 hM3Dq WT; n=17/8 *Mecp2* KO mice; p=0.0023; p=0.9193; Two-way ANOVA). However, chemogenetic inhibition with hM4Di did not affect VSD signals evoked by intracortical stimulation in WT mice (n=16/9, p=0.2099, Two-way ANOVA). VSD signals evoked by intracortical stimulation in mPFC slices from *Mecp2* KO mice expressing hM4Di in mPFC-projecting vHIP neurons were not significantly different than those in mCherry-expressing *Mecp2* KO controls, and were smaller than those in mCherry-expressing WT mice (n=20/10; p=0.9045; p=0.0313; Two-way ANOVA; Figures 4G and 4H). Importantly, there was no significant correlation between performance in the social memory test and the amplitude of VSD signals evoked by intracortical stimulation in hM4Di-expressing *Mecp2* KO mice (n=15, r= −0.07143, p=0.9063, Spearman r correlation), and no difference in these signals between *Mecp2* KO mice that showed improvement in social memory and those that did not (n=11/6 Memory; n=6/4 No memory; p=0.4187; Two-way ANOVA; Figures 4L and 4M). Combined, these results indicate that selective chemogenetic modulation of mPFC-projecting vHIP neurons has specific consequences on the functional strength of this projection, but has smaller effects on other afferent inputs to the mPFC recruited by intracortical stimulation.

We next acutely manipulated neuronal activity to test whether the vHIP-mPFC projection is required for social memory recall, as opposed to the maintenance or initial formation of a social memory. We removed littermate sentinels from the home cage and administered a single intraperitoneal (i.p.) injection of CNO (1 mL/0.5 mg/100 g body weight) two hours before the unrestricted social test (Figure 5A). These i.p. CNO injections did not affect social memory in mCherry-expressing WT mice, whereas acute excitation of mPFC-projecting vHIP neurons by CNO activation of hM3Dq impaired social memory in WT mice (n=8 mCherry WT mice, p=0.0135; n=10 hM3Dq WT mice, p=0.6052, Student’s paired t-test; Figure 5B). Acute inhibition of mPFC-projecting vHIP neurons with the inhibitory DREADD hM4Di in *Mecp2* KO mice caused them to follow the familiar mouse more than the novel mouse (n=10, p=0.0469, Student’s paired t-test; Figure 5B); this effect was opposite to that of chronic inhibition of the vHIP-mPFC projection in *Mecp2* KO mice (see Figure 4C) and the preference of nonsurgical, untreated WT mice to follow the novel mouse (see Figure 2G). Despite these differences in social preference with WT mice, acute inhibition of the vHIP-mPFC projection caused *Mecp2* KO mice to display a significant preference in targeted social interaction, in contrast to control mCherry-expressing *Mecp2* KO mice.

**Figure 5.**
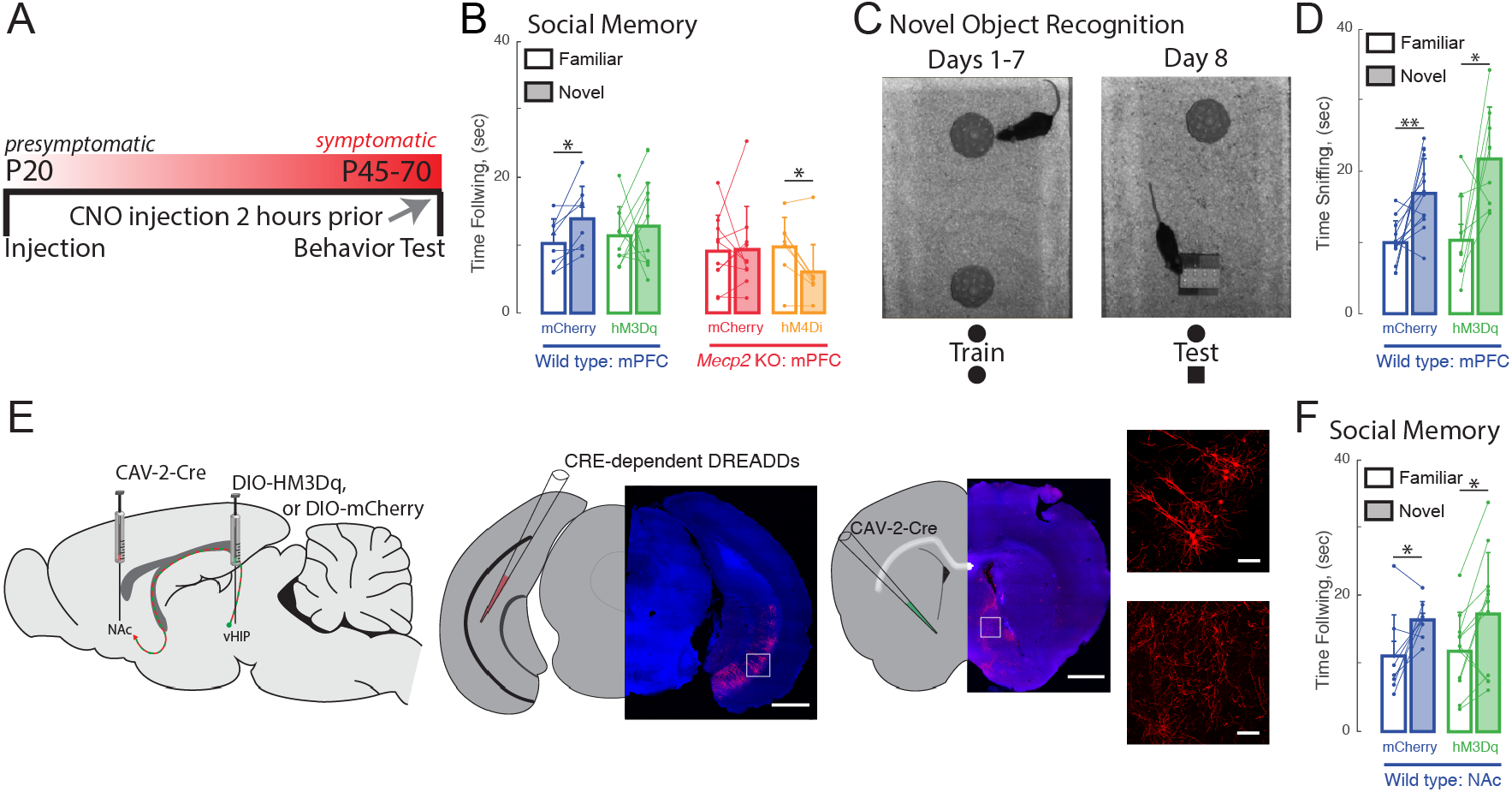
Acute manipulation of activity of vHIP-mPFC projection neurons regulates social memory in a task-and projection-specific manner. **(A)** Experimental timeline for acute DREADD manipulation of the vHIP-mPFC projection. **(B)** Time spent following either familiar or novel mice in unrestricted social interaction, scored by *JAABA* (n = 8 vHIP-mPFC mCherry WT mice, p = 0.0135 Student’s paired t-test; n = 10 vHIP-mPFC hM3Dq WT mice, p = 0.6052 Student’s paired t-test; n = 10 vHIP-mPFC mCherry *Mecp2* KO mice, p = 0.9219 Wilcoxon paired test; n = 8 vHIP-mPFC hM4D1 *Mecp2* KO mice, p = 0.0469 Wilcoxon paired test). **(C)** Schematic of novel object recognition test. **(D)** Time spent sniffing either the familiar or novel object, scored by *JAABA* (n = 8 vHIP-mPFC mCherry WT mice, p = 0.006 Student’s paired t-test; n = 10 vHIP-mPFC hM3Dq WT mice, p = 0.0266 Student’s paired t-test). **(E)** Schematic of CAV2-Cre and DREADD injections to manipulate the vHIP-NAc projection. Injection sites show sparse mCherry labeling of vHIP neurons with identifiable axons in the NAc. Time spent following either familiar or novel mice in unrestricted social interaction, scored by *JAABA* (n = 10 vHIP-NAc mCherry WT mice, p = 0.0391 Wilcoxon paired test; n = 11 vHIP-NAc hM3Dq WT mice, p = 0.0189 Student’s paired t-test). Mean ± SD; * p < 0.05, ** p < 0.01. See also Figure S7.

We next determined whether the vHIP-mPFC projection encodes novelty in general, or specifically social novelty, by testing the acute effect of its activation on novel object recognition. WT mice expressing the excitatory DREADD hM3Dq in mPFC-projecting vHIP neurons showed the same preference for the novel inanimate object as control mCherry-expressing mice two hours after a single i.p. injection of CNO (n=8 WT mCherry mice, p=0.0060; n=10 hM3Dq, p=0.0266; Student’s paired t-test; Figures 5C and 5D), indicating that altering the activity of the long-range vHIP-mPFC projection does not affect hippocampal-dependent novel object recognition.

Do all vHIP projection neurons contribute to social memory, or just those projecting to the mPFC? To address this question, we injected CAV-2-Cre into the nucleus accumbens (NAc) and either control AAV8-DIO-mCherry or AAV8-DIO-hM3Dq into the vHIP of WT mice for selective excitation of NAc-projecting vHIP neurons (Figure 5E). Both hM3Dq-expressing and mCherry-expressing WT mice spent significantly more time following the novel mouse than the familiar mouse (n=10 WT mCherry mice, p=0.0391; n=11 WT hM3Dq mice, p=0.0189; Student’s paired t-test; Figure 5F), indicating that mPFC-projecting vHIP neurons, but not NAc-projecting vHIP neurons, are necessary for the expression of social memory.

### Altered synaptic connectivity of long-range vHIP-mPFC projections in *Mecp2* KO mice

Differences in the spatiotemporal spread of VSD signals evoked by stimulation of the vHIP fiber bundle in mPFC slices between WT and *Mecp2* KO mice could reflect alterations in the innervation pattern of vHIP axons on different postsynaptic cell types in the mPFC. Morphological and electrophysiological recordings *in vivo* and in *ex vivo* slices from rats and WT mice have demonstrated that pyramidal neurons of the ventral CA1 and subiculum form monosynaptic connections with pyramidal neurons in layers 2/3 and 5, as well as with inhibitory interneurons in the PL and infralimbic (IL) regions of the mPFC (Dembrow et al., 2015; Gabbott et al., 2002; Liu and Carter, 2018; Marek et al., 2018). However, a quantitative analysis of the pattern of vHIP innervation onto different postsynaptic cell types was lacking. To identify the first-order postsynaptic neurons innervated by vHIP axons, we injected the trans-synaptic marker wheat germ agglutinin (WGA) into the vHIP (Figures 6A). After 30 hours to allow axonal and trans-synaptic transport in the mPFC (Ruda and Coulter, 1982), we performed immunohistochemistry for WGA and for the neuronal marker NeuN to account for potential WGA injection variability. We identified postsynaptic excitatory neurons by retrograde labeling and classified them based on their axonal projections as pyramidal tract (PT) neurons by injecting FluoroGold (FG) in the dorsal periaqueductal grey (dPAG), or as intratelencephalic (IT) neurons by injecting FG in the contralateral mPFC (c-mPFC) (Figures 6B-D). We identified postsynaptic inhibitory neurons by immunohistochemistry of the markers parvalbumin (PV), calretinin (CAL), and somatostatin (SOM). Regarding the distribution of WGA-positive neurons and neuronal subtypes across the different layers of the mPFC, there were no differences between WT and *Mecp2* KO mice for any of the cell types (Figure S8). In WT mice, the majority of WGA-labeled, NeuN-positive cells were projection pyramidal neurons, with 52% being IT neurons and 37% PT neurons, followed by 4% PV interneurons, 2% CAL interneurons, and 1% SOM interneurons (Figure 6E). The fraction of PT pyramidal neurons was significantly smaller in *Mecp2* KO mice (16%) (n=9 sections from 3 mice for both WT and *Mecp2* KO mice, p=0.0128, Student’s t-test), whereas the fraction of PV interneurons was significantly larger in *Mecp2* KO mice (9%) (p=0.0477, Student’s t-test; Figure 6F). There were no significant differences in the fraction of IT pyramidal neurons (51%), CAL interneurons, (1%), or SOM interneurons (2%) between *Mecp2* KO and WT mice (p=0.8695; p=0.0621; p=0.2590; Student’s t-test; Figure 6F).

**Figure 6.**
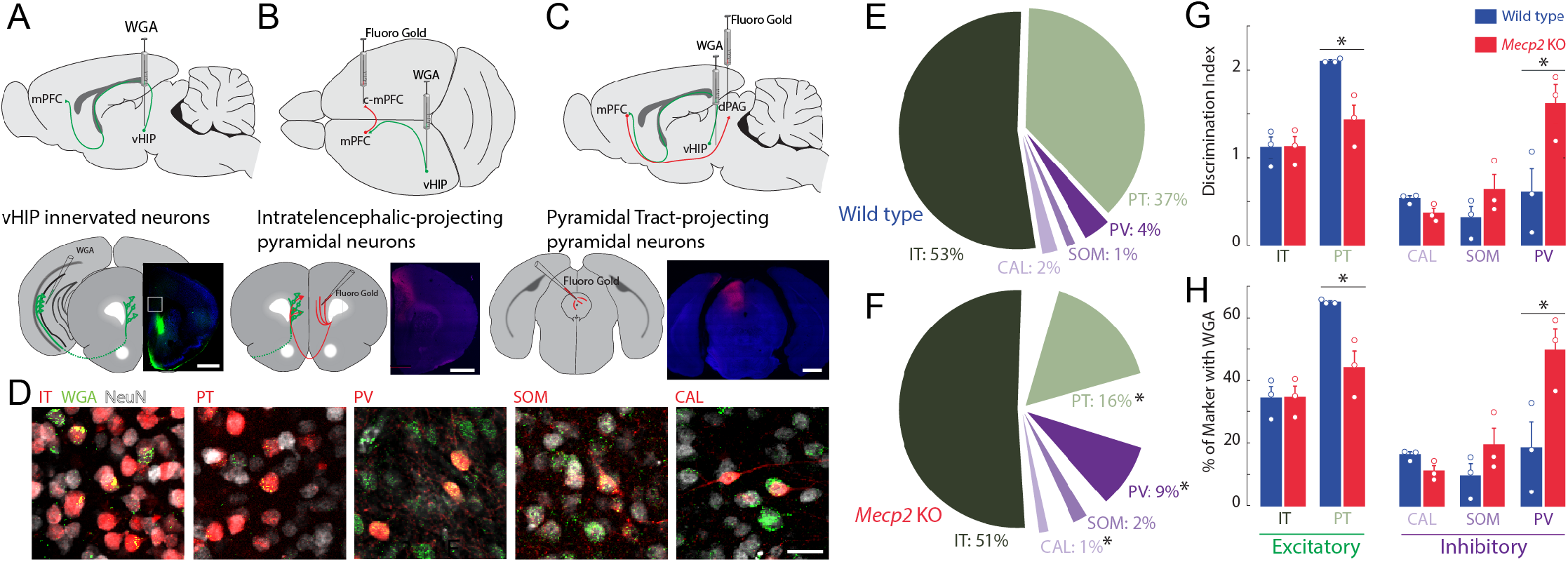
Trans-synaptic tracing of vHIP target neurons in the mPFC. **(A)** WGA injection sites for brains to be used for immunohistochemistry of interneuron markers. Scale bar = 1 mm. **(B)** Injection sites for brains to be used for identifying contralateral projecting mPFC neurons. Scale bar = 1 mm. **(C)** Injection sites for brains to be used for identifying dPAG projecting mPFC neurons. Scale bar = 1 mm. **(D)** Representative examples of WGA identification of inhibitory and excitatory neurons receiving vHIP innervation. Scale bar = 25 μm. **(E-F)** Breakdown of WGA innervated neurons by subtype in WT (**E**) and *Mecp2* KO mice (**F**) (IT p = 0.8695; PT p = 0.0128; CAL p = 0.0621; SOM p = 0.259; PV p = 0.0477, Student’s t-test). **(G)** Discrimination index of innervated cells, with at “at chance” innervation being 1 (IT p = 0.9646; PT p = 0.0161; CAL p = 0.591; SOM p = 0.208; PV p = 0.0425, Student’s t-test). **(H)** Percent of neuron subtype receiving vHIP innervation (IT p = 0.9646; PT p = 0.0161; CAL p = 0.591; SOM p = 0.208; PV p = 0.0425, Student’s t-test) (n = 9 sections 3 mice for all groups). Mean ± SEM; * p < 0.05, ** p < 0.01. See also Figure S8.

Although there is no evidence of neuronal cell death in RTT individuals and *Mecp2*-based mouse models (Chen et al., 2001), we accounted for potential differences in the density of different neuronal cell types by implementing a discrimination index for each mPFC neuron type that was trans-synaptically labeled with WGA. We gave an index of 1 when vHIP axons innervated neurons at chance values (having the same proportion of postsynaptic cell type in NeuN+ and WGA+ populations); by contrast, an index higher than 1 reflected innervation higher than chance and lower than 1 reflected innervation lower than chance. In the mPFC of WT mice, PT pyramidal neurons were preferentially innervated by vHIP axons (index=2.1104, n=9 sections from 3 mice, p=0.0001, one-sample Student’s t-test; Figure 6G). IT pyramidal neurons and PV-positive interneurons did not have a significant discrimination index (index=1.1192 and 0.6050, n=9 / 3 mice each, p=0.4186 and p=0.2754, one-sample Student’s t-test). Further, vHIP innervation of CAL and SOM interneurons occurred with a probability lower than chance (index=0.5346 and 0.3165, n=9/3 each mouse, p=0.0304 and p=0.0322, one-sample Student’s t-test). By contrast, vHIP axons innervated PV cells in the mPFC of *Mepc2* KO mice more than in WT mice (n=9/3, p=0.0425, Student’s t-test), to the detriment of PT pyramidal cells (n=9/3, p=0.0161, Student’s t-test). We observed similar results when we assessed the percent of each neuronal subtype that received vHIP innervation; PV interneurons were innervated more at the expense of PT pyramidal cells in *Mecp2* KO compared to WT mice (Figure 6H). Combined, these results indicate that the pattern of innervation of vHIP axons in the mPFC changes from mainly targeting excitatory projection pyramidal neurons in WT mice to preferentially targeting PV-expressing inhibitory GABAergic interneurons (Figures 6G and 6H).

### Enhanced vHIP-mPFC synaptic strength in *Mecp2* KO mice

Larger peak VSD signals evoked by stimulation of the vHIP fiber bundle in mPFC slices from *Mecp2* KO mice could reflect either more or stronger excitatory synapses between presynaptic vHIP axons and postsynaptic mPFC neurons. We estimated the numerical density and size of *en passant* presynaptic terminals along afferent axons within the mPFC by labeling them with mCherry delivered by AAV2 injected either into the vHIP or the c-mPFC, and then performing immunohistochemistry of the presynaptic vesicle marker VGLUT1 (Figures 7A-E). We performed automated detection and size analysis using *Bouton Analyzer* (Gala et al., 2017) (Figure 7E). The numerical densities of mCherry-expressing *en passant* presynaptic terminals belonging to vHIP neurons and those belonging to c-mPFC neurons were comparable between WT and *Mecp2* KO mice across all cortical layers of the mPFC (p>0.05, One-way ANOVA with Tukey’s Multiple Comparisons; Figure 7F). However, the weighted size of individual presynaptic boutons of vHIP axons in layer 5 of the mPFC was significantly larger in *Mecp2* KO mice (n=993 boutons in WT mice, n=682 *Mecp2* KO mice, p=0.0025, Kolmogorov-Smirnov test), and those in layer 2/3 showed a larger fraction of both small and large boutons than in WT mice (n=543 boutons in WT mice, n=733 in *Mecp2* KO mice, p=4.36 x 10^-05^, Kolmogorov-Smirnov test; Figures 7G and 7H). In contrast, the sizes of presynaptic boutons of c-mPFC axons in layers 5 and 2/3 of the mPFC were significantly smaller in *Mecp2* KO mice (p<0.01; Figures 7G and 7H). Interestingly, presynaptic boutons of vHIP axons in layer 5 of the mPFC were significantly smaller than those of c-mPFC axons in WT mice, a difference absent in *Mecp2* KO mice due to larger vHIP boutons and smaller c-mPFC boutons (Figure 7G). These results are reminiscent of the amplitude of VSD signals in *ex vivo* mPFC slices evoked by either vHIP or intracortical stimulation, which were biased towards intracortical stimulation in WT mice, and of comparable amplitude in *Mecp2* KO mice due to both larger vHIP-evoked VSD signals and smaller intracortical-evoked responses (see Figure 3L).

**Figure 7.**
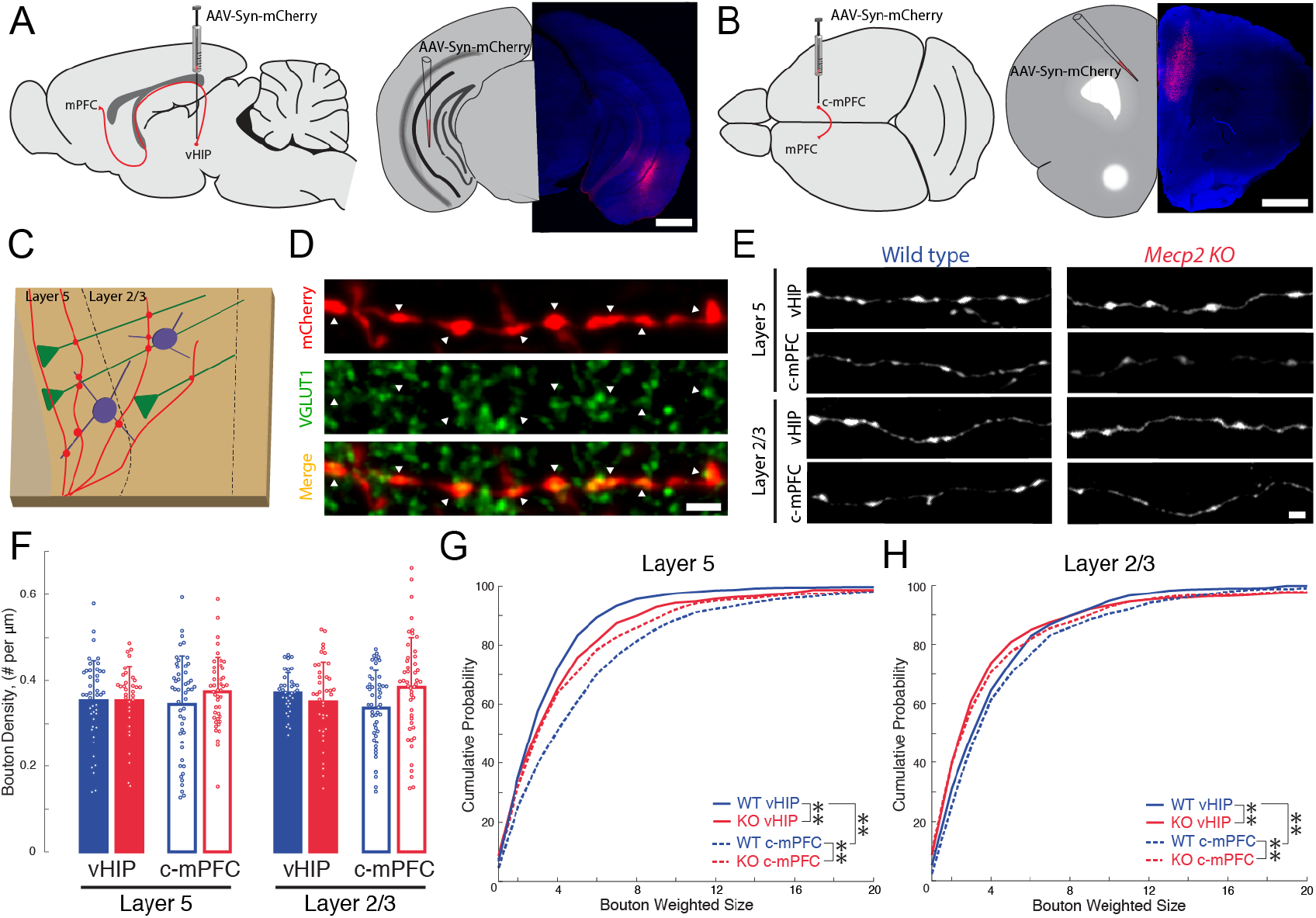
The size of presynaptic boutons is altered in *Mecp2* KO mice. **(A-B)** Schematic and representative examples of AAV2-hSyn-mCherry injection sites for identifying vHIP axons (**A**) or contralateral mPFC axons (**B**). Scale bars 1 mm. **(C)** Schematic of vHIP axons in the mPFC. **(D)** Axonal swellings identified as presynaptic boutons co-labeled for mCherry and VGLUT1. Scale bar 2 μm **(E)** Representative examples of mCherry-filled presynaptic boutons. Scale bar 2 μm. **(F)** Numerical densities of axonal boutons per length of vHIP and c-mPFC axons located in layer 5 or 2/3 of mPFC (n=45 axons WT L5 vHIP; n=36 *Mecp2* KO L5 vHIP; n=49 WT L5 c-mPFC; n=43 KO L5 c-mPFC; n=37 WT L2/3 vHIP; n=41 KO L2/3 vHIP; n=49 WT L2/3 c-mPFC; n=49 KO L2/3 c-mPFC; WT L5 vHIP vs. *Mecp2* KO L5 vHIP, p>0.9999; WT L5 vHIP vs. WT L5 c-mPFC, p=0.9999; *Mecp2* KO L5 vHIP vs. *Mecp2* KO L5 c-mPFC, p=0.9779; WT L5 c-mPFC vs. *Mecp2* KO L5 c-mPFC, p=0.8218; WT L2/3 vHIP vs. *Mecp2* KO L2/3 vHIP, p=0.9799; WT L2/3 vHIP vs. WT L2/3 c-mPFC, p=0.6790; *Mecp2* KO L2/3 vHIP vs. *Mecp2* KO L2/3 c-mPFC, p=0.6914; WT L2/3 c-mPFC vs. *Mecp2* KO L2/3 c-mPFC, p=0.1797; One-way ANOVA with Tukey’s multiple comparisons). Mean ± SD. **(G)** Cumulative probability distributions of the estimated size of presynaptic boutons comparing vHIP and c-mPFC axons in mPFC layer 5 of WT and *Mecp2* KO mice (vHIP boutons in layer 5 of WT mice vs. *Mecp2* KO mice, p=0.0025; c-mPFC boutons in layer 5 of WT mice vs. *Mecp2* KO mice, p=1.31×10^-5^; vHIP vs. c-mPFC boutons in layer 5 of WT mice, p=2.93×10^-20^; vHIP vs. c-mPFC boutons in layer 5 of *Mecp2* KO mice, p=0.2727; n=993 WT L5 vHIP boutons; n=682 KO L5 vHIP; n=792 WT L5 c-mPFC; n=577 KO L5 c-mPFC; Kolmogorov-Smirnov test). **(H)** Cumulative probability distributions of bouton sizes comparing vHIP and c-mPFC axons in layer 2/3 of WT and *Mecp2* KO mice (vHIP boutons in layer 2/3 of WT mice vs. *Mecp2* KO mice, p=4.36×10^-5^; c-mPFC boutons in layer 2/3 of WT mice vs. *Mecp2* KO mice, p=9.72×10^-9^; vHIP vs. c-mPFC boutons in layer 2/3 of WT mice, p=0.0151; vHIP vs. c-mPFC boutons in layer 2/3 of *Mecp2* KO mice,

Because the size of presynaptic terminals is positively correlated with presynaptic vesicle density (Harris and Weinberg, 2012), which in turn is positively correlated with synaptic strength (Murthy et al., 2001), we reasoned that larger VSD responses to vHIP fiber stimulation could reflect enhanced synaptic strength. To selectively stimulate different axonal projections onto the same postsynaptic mPFC neuron during whole-cell intracellular recordings, we used two opsins with shifted excitation spectra (Klapoetke et al., 2014). We injected AAV2s expressing the red-shifted opsin Chrimson into the ipsilateral vHIP, and those expressing the blue-shifted opsin Chronos into the c-mPFC (Figures 8A, 8B, S9A, and S9B). In mice expressing Chrimson in the vHIP and Chronos in the c-mPFC, a brief (1-4 ms) pulse of either red (630nm) or blue (430nm) light evoked monotonic inward currents in layer 5 neurons in the presence of 4-AP (100mM), TTX (1μM), and 4mM Ca^2+^, which represent monosynaptic excitatory postsynaptic currents (EPSCs) (Petreanu et al., 2009). The amplitude of red light vHIP-evoked EPSCs was significantly larger in pyramidal neurons from *Mecp2* KO mice compared to those from WT mice (n=15 cells in 5 slices from 5 WT mice; n=11/5/5 *Mecp2* KO mice; p=0.0190; Two-way ANOVA; Figure 8C), whereas the amplitude of vHIP-evoked monosynaptic EPSCs in layer 5 putative interneurons (identified by their size, shape, input resistance, and capacitance; see Figures S9C and S9D) was comparable in both genotypes (n=6 cells in 4 slices from 4 WT mice; n=6/6/6 *Mecp2* KO mice; p=0.3206; Two-way ANOVA; Figure 8D). By contrast, blue light stimulation of Chronos-expressing c-mPFC axons evoked EPSCs of comparable amplitude in both layer 5 pyramidal neurons and interneurons from WT and *Mecp2* KO mice (p=0.1446; p=0.9685; Two-way ANOVA; Figures 8E and 8F). The cell-by-cell normalization of the amplitude of EPSCs evoked by red light stimulation of Chrimson-expressing vHIP axons to the amplitude of EPSCs evoked by blue light stimulation of Chronos-expressing c-mPFC axons revealed that vHIP-evoked EPSCs were larger in layer 5 pyramidal from *Mecp2* KO mice compared to WT mice (103% vs. 54%; p=0.0008; Two-way ANOVA) (Figure 8G). By contrast, normalized EPSC amplitudes in putative interneurons were not significantly different between *Mecp2* KO and WT mice (88% vs. 67%; p=0.0604; Two-way ANOVA) (Figure 8H). These results are reminiscent of the normalized amplitude of VSD signals evoked by stimulation of the fluorescently labeled vHIP fiber bundle compared to intracortical stimulation (see Figure 3L).

**Figure 8.**
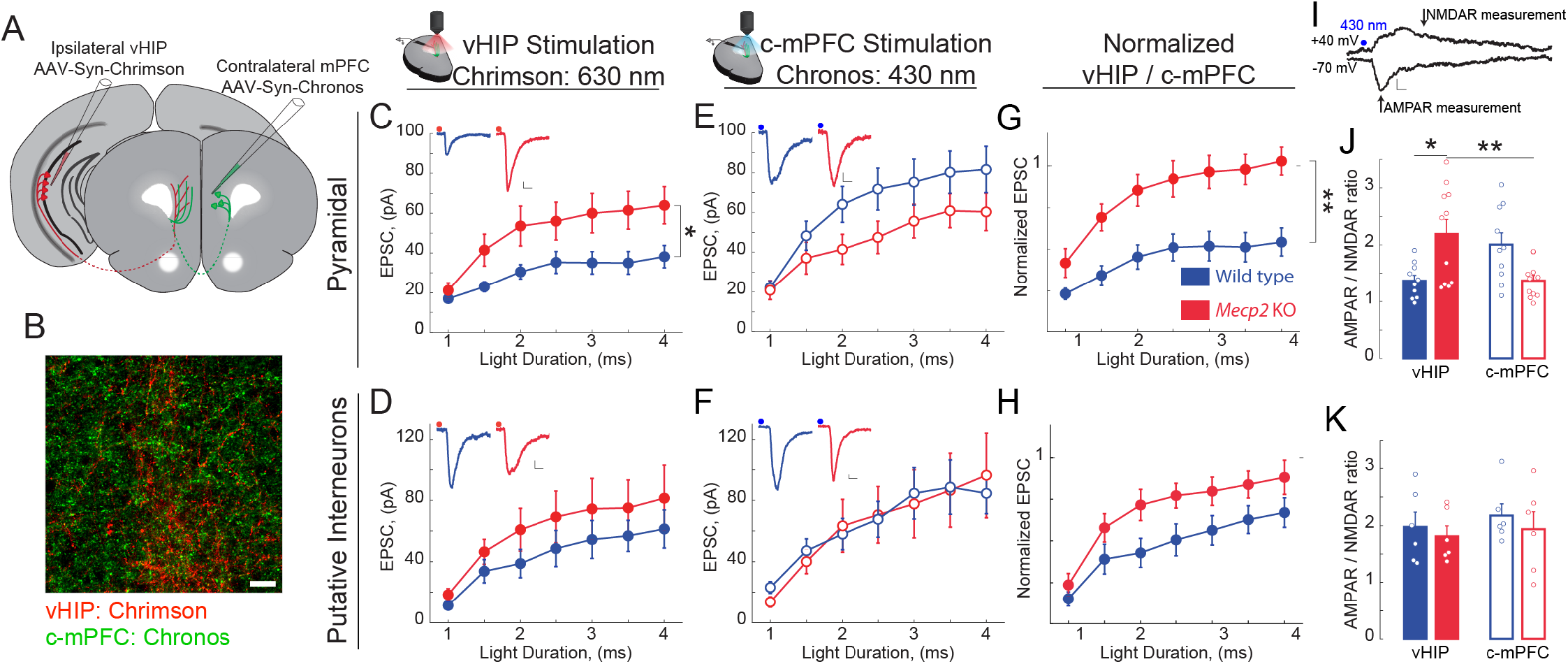
vHIP synapses on mPFC layer 5 pyramidal neurons are stronger in *Mecp2* KO mice. **(A)** Schematic of injection sites for Chrimson in the ipsilateral vHIP and Chronos in the contralateral mPFC. **(B)** Representative image of an mPFC slice with Chrimson-expressing vHIP afferents and Chronos-expressing c-mPFC afferents. Scale bar 50 μm. **(C & D)** Input-output relationship of vHIP afferent (red light) evoked responses in pyramidal neurons (**C** p=0.0190, Two-way ANOVA) and putative interneurons (**D**, p=0.3206, Two-way ANOVA) with representative traces (inset). **(E & F)** Input-output relationship of c-mPFC afferent (blue light) evoked responses in pyramidal neurons (**E**, p=0.1446, Two-way ANOVA) and putative interneurons (**F**, p=0.9685, Two-way ANOVA) with representative traces (inset) (Scale bars 10 pA/12 ms). **(G & H)** The amplitude of vHIP afferent (red light)-evoked EPSCs was normalized to the peak EPSCs evoked by blue light stimulation of c-mPFC afferents in pyramidal neurons (**G**, p=0.0008, Two-way ANOVA) and putative interneurons (**H**, p=0.0604, Two-way ANOVA) (n=15 cells from 5 slices from 5 mice WT pyramidal neurons, 6/4 WT putative interneurons, 11/5 KO pyramidal cells, 6/6/6 KO putative interneurons). **(I)** Representative example trace for the measurement of AMPAR/NMDAR ratio (Scale bar 5 pA/10 ms). **(J & K)** AMPAR/NMDAR ratios of vHIP afferent (red light)-evoked and c-mPFC afferent (blue light)-evoked responses in pyramidal cells (**J**, WT vs. *Mecp2* KO vHIP, p=0.0114, WT vs. *Mecp2* KO c-mPFC, p=0.0727; WT vHIP vs. WT c-mPFC, p=0.0809; *Mecp2* KO vHIP vs. *Mecp2* KO c-mPFC, p=0.0094; One-way ANOVA with Tukey’s multiple comparisons) and putative interneurons (**K**, WT vs. *Mecp2* KO vHIP, p=0.9620; WT vs. *Mecp2* KO c-mPFC, p=0.8976; WT vHIP vs. WT c-mPFC, p=0.9369; *Mecp2* KO vHIP vs. *Mecp2* KO c-mPFC, p=0.9825; One-way ANOVA with Tukey’s multiple comparisons) (n=10 cells from 5 slices from 5 mice WT pyramidal cells; 6/4/4 WT putative interneurons; 10/5/5 *Mecp2* KO pyramidal cells for both vHIP and c-mPFC stimulation; 6/6/6 *Mecp2* KO putative interneurons). Mean ± SEM; * p<0.05, ** p<0.01. See also Figure S9.

As a direct measure of synaptic strength, we calculated the ratio of the α-amino-3-hydroxy-5-methyl-4-isoxazolepropionic acid (AMPA) receptor component of the EPSC (recorded at −70 mV) to that of the NMDA receptor component (recorded at +40 mV) in the same neuron (Figure 8I). The AMPAR/ NMDAR ratio of EPSCs evoked by red light vHIP stimulation was larger in layer 5 pyramidal neurons from *Mecp2* KO mice compared to those from WT mice (n=10 cells from 5 WT mice; n=11/5 *Mecp2* KO mice; p=0.0114; One-way ANOVA with Tukey’s Multiple Comparisons), whereas the AMPAR/NMDAR ratio of EPSCs evoked by blue light c-mPFC stimulation was not different in *Mecp2* KO mice compared to WT mice (p=0.0727; One-way ANOVA with Tukey’s Multiple Comparisons; Figure 8J). By contrast, the AMPAR/NMDAR ratio of EPSCs evoked by either vHIP or c-mPFC stimulation in putative interneurons was not significantly different between *Mecp2* KO and WT mice (n=6 cells from 4 WT mice; n=6/6 *Mecp2* KO mice; p=0.9620 vHIP stimulation; p=0.8976 c-mPFC stimulation; One-way ANOVA with Tukey’s Multiple Comparisons; Figure 8 K). Combined, these results indicate that vHIP excitatory synapses are selectively stronger on layer 5 pyramidal neurons, but not interneurons, in the mPFC of *Mecp2* KO mice.

## DISCUSSION

Here, we characterized the projection from the vHIP to the PL region of the mPFC at structural and functional levels in WT mice, and described its atypical features in the *Mecp2* KO model of Rett syndrome. Because the vHIP has been implicated in social memory, and due to the involvement of the mPFC PL subregion in sociability and social novelty encoding, we tested the role of the vHIP projection to the mPFC in social behaviors. By chemogenetically manipulating neuronal activity selectively in mPFC-projecting vHIP CA1 neurons, we demonstrated that these projection neurons regulate social memory in a specific and selective manner, only influencing the discrimination between social targets without affecting other aspects of social interactions.

Coherent, synchronous oscillations between the vHIP and the mPFC underlie working memory tasks in rats (Gordon, 2011). Entrainment of these oscillations occurs in the first postnatal week (Brockmann et al., 2013) and is driven by monosynaptic glutamatergic projections from pyramidal neurons of the ventral CA1 and subiculum to pyramidal neurons and GABAergic interneurons in the IL and PL subregions of the mPFC (Anastasiades et al., 2018; Dégenètais et al., 2003; Dembrow et al., 2010; Thierry et al., 2000). Despite the wealth of information linking this projection to neuropsychiatric disorders (Li et al., 2015), little is known about the synaptic and cellular bases of this long-range projection. To better define the connectivity of this circuit, we performed trans-synaptic tracing to identify postsynaptic targets of vHIP afferents and determined how this innervation pattern is altered in *Mecp2* KO mice. Although recent evidence suggests that monosynaptic excitatory vHIP inputs to the mPFC are strongest on pyramidal neurons in layer 2/3 of the IL, but dominated by feed-forward inhibition onto them (Marek et al., 2018), there is also evidence of strong monosynaptic excitatory innervation of IT-projecting pyramidal neurons in layer 5 of the PL (Liu and Carter, 2018). Our trans-synaptic tracing data support the latter observations, with over 50% of WGA-positive neurons being IT-projecting pyramidal neurons. We also identified PV-positive interneurons as the inhibitory subgroup most innervated by vHIP axons, which is consistent with the observation of vHIP-driven feed-forward inhibition in the IL (Marek et al., 2018). In *Mecp2* KO mice, there are fewer vHIP-innervated PT-projecting pyramidal neurons, which have been shown to encode social dominance (Franklin et al., 2016). This suggests a basis for the impairments in social memory performance observed in *Mecp2* KO mice. However, there are more vHIP-innervated PV-positive interneurons, which may lead to tonic inhibition of the mPFC network due to the hyperactivity of the vHIP in *Mecp2* KO mice (Calfa et al., 2011). Interestingly, the density of *en passant* presynaptic boutons along vHIP axons in the mPFC was not altered in *Mecp2* KO mice, indicating a redistribution of excitatory vHIP inputs on different cell types in the mPFC, which results in an atypical wiring pattern.

Although the number of vHIP boutons in the mPFC was not altered, their individual volume was larger in layer 5 of the mPFC of *Mecp2* KO mice. By contrast, vHIP boutons in layer 2/3 had a more prominent bimodal distribution of volumes compared to WT mice. Because the size of presynaptic boutons is correlated with synaptic strength (Murthy et al., 2001), we tested the strength of vHIP-mPFC synapses onto both pyramidal neurons and putative interneurons. EPSCs evoked by optogenetic excitation of vHIP fibers in layer 5 mPFC pyramidal cells were larger and had higher AMPAR/NMDAR ratios in *Mecp2* KO mice compared to WT littermates. Interestingly, vHIP-evoked EPSC amplitudes and AMPAR/NMDAR ratios in putative interneurons were not affected in *Mecp2* KO mice. Together with the trans-synaptic identification of postsynaptic targets of vHIP terminals in the mPFC, these results indicate that vHIP axons in *Mecp2* KO mice innervate fewer pyramidal neurons with stronger synaptic strength, but they have similar innervation strength onto more inhibitory interneurons. Such altered connectivity is reflected in the pattern of vHIP-evoked neuronal depolarizations in mPFC slices revealed by high-speed voltage imaging. In these studies, the larger amplitudes reflect stronger excitatory synapses onto pyramidal neurons, and the spatiotemporal spread throughout the mPFC slice and over time reflects altered connectivity and feed-forward inhibition. It is possible that a larger proportion of inhibitory neurons is chronically activated by hyperactive vHIP inputs in *Mecp2* KO mice, causing tonic inhibition of the mPFC network, and that vHIP activation of IT-projecting pyramidal neurons is able to overcome this inhibition.

Deficits in the E/I balance within the mPFC have been linked to impaired sociability in WT mice, as well as in mouse models of ASDs (Selimbeyoglu et al., 2017; Yizhar et al., 2011; Brumback et al., 2017). In addition, the hippocampal network is integral to the expression of social memory (Hitti and Siegelbaum, 2014; Meira et al., 2018; Okuyama et al., 2016). Because network activity within the vHIP and the mPFC, as well as the projection pattern of vHIP afferents in the mPFC, are altered in *Mecp2* KO mice, we characterized their sociability, social interaction, and social memory. We performed these experiments using computer vision to track multiple freely interacting mice and a computer learning algorithm trained to identify different behaviors (Kabra et al., 2013; Ohayon et al., 2013; Robie et al., 2017). This automatic and unbiased screen of social interactions revealed that, despite showing typical sociability, *Mecp2* KO mice displayed an atypical behavior of jumping at other mice more than following or sniffing, which was not associated with aggression. This finding, to our knowledge, is the first description of an atypical social interaction in a mouse model of ASD. We also identified a deficit of social memory in *Mecp2* KO mice, which failed to discriminate between a co-housed littermate and a novel mouse as the target of their social interactions under unrestricted conditions.

To demonstrate a causal role of altered vHIP-mPFC inputs on atypical social behaviors in *Mecp2* KO mice, we mimicked the characteristic hippocampal hyperactivity of *Mecp2* KO mice in WT mice by chronic chemogenetic activation with the excitatory hM3Dq DREADD selectively expressed in mPFC-projecting vHIP neurons. Such chronic excitation impaired social memory, without affecting other types of social interactions. Chronic inhibition of vHIP-mPFC projecting neurons in WT mice also impaired social memory, indicating that there is a set level of proper neuronal activity in this long-range projection, with any deviation resulting in social memory deficits. Even though the vHIP-mPFC projection has been causally tied to anxiety-like behaviors (Padilla-Coreano et al., 2016), chronic chemogenetic manipulation of vHIP-mPFC projection neurons had no major consequences on the time spent in the center of the arena, the time spent engaging in grooming, or the overall locomotor behavior, compared to CNO-treated WT mice expressing Cre-driven mCherry in mPFC-projecting vHIP neurons. This finding demonstrates the selectivity of this manipulation of the vHIP-mPFC projection to social memory. Voltage imaging of network responses in mPFC slices from hM3Dq-expressing mice revealed stronger vHIP inputs and, surprisingly, weaker responses evoked by stimulation of intracortical inputs (which were not chemogenitically manipulated), resembling the responses observed in *Mecp2* KO mice. These results may reflect feed-forward homeostatic mechanisms within the mPFC microcircuit, as has been hypothesized to give rise to the dichotomy in the direction of E/I imbalances between limbic and cortical structures in models of neuropsychiatric diseases (Nelson and Valakh, 2015).

Despite the fact that many brain regions are dysfunctional in *Mecp2* KO mice, selective chronic inhibition of vHIP-mPFC projection neurons was sufficient to improve their social memory. In addition, social memory discrimination scores were negatively correlated with the amplitude of vHIP-driven depolarizations in mPFC slices: larger voltage dye signals corresponded to worse social memory performance, whereas smaller voltage dye signals corresponded with better expression of social memory. By contrast, chronic inhibition of vHIP-mPFC projection neurons did not affect voltage dye responses evoked by intracortical stimulation, and these responses did not correlate with social memory performance.

Acute manipulation of vHIP-mPFC projection neurons immediately prior to the behavioral test also affected social memory: their excitation with hM3Dq impaired social memory in WT mice, whereas inhibition with hM4Di improved social discrimination in *Mecp2* KO mice. These data suggest that the vHIP-mPFC projection is necessary for the recall of social memory, as opposed to its maintenance or initial formation. Because acute manipulation impaired social memory in WT mice, we used this paradigm to test the specificity of the vHIP-mPFC projection to the social aspect of memory, as well as its selectivity by testing another vHIP projection target. Acute excitation of hM3Dq-expressing mPFC-projecting vHIP neurons did not affect novel object recognition in WT mice, indication that this manipulation did not affect overall hippocampal function. Furthermore, regulation of social memory was selective to mPFC-projecting vHIP neurons because excitation of hM3Dq-expressing vHIP neurons projecting to the NAc did not affect social memory, which is at odds with a previous report concluding that vHIP-NAc projection is required for social memory (Okuyama et al., 2016). A key difference between these studies is our intersectional viral approach to selectively and exclusively express DREADDs in specific projection neurons of the vHIP.

In summary, we demonstrate that the vHIP-mPFC projection regulates social memory in WT mice, and that its dysfunction causes social memory deficits in *Mecp2* KO mice. Defining the synaptic bases of social behaviors provides insight and potential targets for therapies in psychiatric disorders associated with vHIP-mPFC dysfunction, such as autism and schizophrenia.

## Acknowledgements

This research was supported by NIH grant R56-NS103089-01A1 (LP-M), the Civitan International Foundation (MLP), NIH training grant T32NS061788-04 (MLP), and *Rettsyndrome.org* (LP-M). We thank Dr. Wei Li for comments on the manuscript, Ms. Karen Ayala-Baylon for mouse colony maintenance, and Dr. Takafumi Inoue (Waseda University, Tokyo, Japan) for electrophysiology and imaging software. We also thank Mr. Rahul Gaini, Ms. Allison Fusilier, and Ms. Melissa Bentley for their contributions to data acquisition and analysis in early phases of this project.

## Author Contributions

M.L.P. and L.P-M. contributed to the study design. M.L.P. and H.A.R. contributed to data collection. M.L.P. and H.A.R. conducted surgeries, behavioral experiments, and histological analyses. M.L.P. performed electrophysiological experiments and analyses for all experiments. M.L.P. and L.P-M. wrote the paper. All authors discussed and commented on the manuscript.

## Declaration of Interest

The authors declare no competing financial interests.

## KEY RESOURCES TABLE

| REAGENT or RESOURCE | SOURCE | IDENTIFIER |
| --- | --- | --- |
| **Antibodies** |  |  |
| Anti-mCherry | Abcam | ab167453 |
| Anti-GFP | Abcam | ab13970 |
| Anti-PV | Abcam | ab11427 |
| Anti-PV | Synaptic Systems | 195004 |
| Anti-cFos | Synaptic Systems | 226004 |
| Anti-CAL | Synaptic Systems | 214106 |
| Anti-SST | Synaptic Systems | 366004 |
| Anti-NeuN | Synaptic Systems | 266004 |
| Anti-Lectin (WGA) | Sigma | T4144-1VL |
| Alexa Fluor 647 Goat Anti-Mouse | Jackson ImmunoResearch | 115-605-003 |
| Biotinylated Goat Anti-Rabbit | Vector Laboratories | BA-1000 |
| Biotinylated Goat Anti-Guinea Pig | Vector Laboratories | BA-7000 |
| Streptavidin, Alexa Fluor 405 | Thermo Fisher Scientific | S-32351 |
| Streptavidin, Alexa Fluor 488 | Thermo Fisher Scientific | S-32354 |
| Streptavidin, Alexa Fluor 594 | Thermo Fisher Scientific | S-32356 |
| **Bacterial and Virus Strains** |  |  |
| AAV2-hSyn-Chronos-GFP | UNC Vector Core |  |
| AAV2-hSyn-Chrimson-tdTomato | UNC Vector Core |  |
| AAV2-hSyn-mCherry | UNC Vector Core |  |
| CAV-2-Cre-GFP | CNRS Biocampus Montpellier |  |
| AAV8-DIO-hSyn-hM4Di-mCherry | Addgene |  |
| AAV8-DIO-hSyn-hM3Dq-mCherry | Addgene |  |
| AAV8-DIO-hSyn-mCherry | Addgene |  |
| **Chemicals, Peptides, and Recombinant Proteins** |  |  |
| Dextran-Alexa-594 10,000 MW | Thermo Fisher Scientific | D22913 |
| Wheat Germ Agglutinin | Vector Laboratories | L-1020 |
| FluoroGold | Fluorochrome |  |
| Retrobeads | Lumafluor Inc. |  |
| Tetrodotoxin | Alomone Labs |  |
| 4-Aminopyridine | Sigma | 275875 |
| D-AP5 | Tocris | 106 |
| **Experimental Models: Organisms/Strains** |  |  |
| Mouse: *Mecp2*<sup>tm1.1Jae</sup> | Mutant Mouse Resource and Research Center | MMRRC:000415-UCD |
| **Software and Algorithms** |  |  |
| *Motr* | Janelia Research Campus |  |
| *JAABA* | Janelia Research Campus |  |
| **Other** |  |  |
| Born Blonde Maxi | Clairol |  |

## CONTACT FOR REAGENT AND RESOURCE SHARING

Further information and requests for resources and reagents should be directed to and will be fulfilled by the Lead Contact, Lucas Pozzo-Miller.

## EXPERIMENTAL MODEL AND SUBJECT DETAILS

Female mice with deletions of exon 3 in the *Mecp2* gene (Mecp2^tm1.1Jae^; Chen et al., 2001) were obtained from the Mutant Mouse Regional Resource Center (University of California, Davis), and maintained in a pure C57/ BL6 background by crossing them with male WT C57/BL6 mice. All experimental subjects were male hemizygous Mecp2^tm1.1Jae^ mice, referred to as KO mice. Subjects classified as presymptomatic were tested between P20 and P24. Subjects that exhibited Rett-like symptoms, such as hypoactivity, hind limb clasping, resting tremors, and reflex impairments (Guy et al., 2001), were classified as symptomatic (between P45-P60). Age-matched male WT littermates were used as controls. Mice were handled and housed according to the Committee on Laboratory Animal Resources of the National Institutes of Health. All experimental protocols were reviewed and approved annually by the Institutional Animals Care and Use Committee of the University of Alabama at Birmingham.

## METHOD DETAILS

### Intracranial injections

Mice were anesthetized with 4% isoflurane vapor in 100% oxygen gas and maintained with 1-2.5% isoflurane vapor in 100% oxygen gas mixtures. Mice were aligned in a stereotactic frame (Kopf Instruments, Tujunga, CA), and their body temperature was measured with a rectal probe and maintained with a heating pad. Midline incisions were was made down the scalps, and a dental drill performed craniotomies. A 2.5 μL syringe (Hamilton Company, Reno, NV) delivered the injection solutions (viruses, RetroBeads, fluorescent dextran, FluoroGold, WGA) at a rate of 0.25 μL/min using a microsyringe pump (UMP3 UltraMicroPump, Micro4; World Precision Instruments, Sarasota, FL). The needle was slowly extracted from the injection site over 10 min, after which the incision was closed with surgical glue. All ages and coordinates for each experiment are relative to the bregma and listed as anterior/posterior (A/P), medial/lateral (M/L), and dorsal/ventral (D/V).

### Immunohistochemistry

Mice were anesthetized with an i.p. injection of ketamine (100 mg/kg) and transcardially perfused with ice-cold 1X phosphate-buffered saline (PBS), followed by ice-cold 4% paraformaldehyde (PFA) in 1X PBS. The brain was dissected and postfixed in 4% PFA overnight. Brains were sectioned at 30 μm thickness with a vibratome (PELCO 100, model 3000; Ted Pella Inc., Redding, CA) and stored at 4°C in 1X PBS containing 0.01% sodium azide. Free-floating sections were permeabilized using 0.25% Triton-100X for 15 min and subsequently incubated in blocking solution (0.01% sodium azide, 2% bovine serum albumin (BSA), 0.1% Triton-100X, 2M glycine, and 10% goat serum in 1X PBS) for 1 h. Antibody diluent consisted of 0.01% sodium azide, 2% BSA, 0.1% Triton-100X, and 5% goat serum in 1X PBS. Primary antibodies were diluted in antibody diluent at concentrations listed below and incubated for 36 h at room temperature. After washing three times for 5 min in 1X PBS, secondary antibodies were diluted in antibody diluent and incubated for 4 h at room temperature. Sections were washed three times for 5 min in 1X PBS before mounting with Vectashield mounting media (Vector Biolabs, Malvern, PA).

#### Amplification

For primary antibodies requiring further amplification (c-Fos, mCherry, and GFP), a biotinylation step was added following the primary antibody incubation. Sections were incubated in biotinylated anti-host of the primary antibody (Vector Biolabs, Malvern, PA) at a concentration of 1:200 in antibody diluent for 2 h. After washing three times for 5 min in 1X PBS, sections were incubated in streptavidin-conjugated fluorophore (Alexa-405, Alexa-488, Alexa-594; Thermo Fisher Scientific, Waltham, MA) diluted in antibody diluent at a concentration of 1:1600 for 4 h. Sections were washed three times for 5 min before mounting with Vectashield mounting media (Vector Biolabs, Malvern, PA).

#### Injection of WGA

For identification of mPFC neurons innervated by the vHIP, P45 mice were injected with 500 nL of 4% WGA (Vector Laboratories, Burlingame, CA) (Ruda and Coulter, 1982) into the vHIP (3.8 A/P, 3.3 M/L, 3.5 D/V from the bregma). Mice were sacrificed 30 h later, a time point that our pilot screen confirmed would enhance the monosynaptic transfer labelling of primary neurons. To avoid cross-reactivity between the anti-WGA primary antibody and other antibodies, sections were first incubated with anti-WGA and then underwent subsequent biotinylation and streptavidin steps. Following the last wash, sections were again blocked for 1 h. Immunohistochemitry for neuronal subtypes and NeuN was subsequently performed following the standard protocol.

#### Injection of FluoroGold

For identification of mPFC pyramidal neurons based on their projections, P30 mice were injected with 250 nL of 2% FluoroGold (Fluorochrome, Denver, CO) (Schmued and Fallon, 1986) in either the right mPFC (1.45 A/P, 0.5 M/L, 1.45 D/V from the bregma) or the dPAG (1.18 A/P, 4.2 M/L, 2.36 D/V from the bregma with a needle angel of 26∘). At P45, the same mice were injected with 500 nL of 4% WGA into the vHIP (3.8 A/P, 3.3 M/L, 3.5 D/V from the bregma) and sacrificed 30 h later.

#### Identification of socially-activated neurons

For identification of vHIP projection neurons, P35 mice were injected with red RetroBeads (Lumafluor, Durham, NC) (Quattrochi et al., 1989) in the LH (1.34 A/P, 1.1 M/L, 5.3 D/V from the bregma) and green RetroBeads in the mPFC (1.45 A/P, 0.5 M/L, 1.45 D/V from the bregma). Two weeks after surgery, mice underwent 3 days of testing acclimation (3 min of handling and 10 min inside the 16×10 in test box. On day 4, mice were placed in the test box containing either a littermate sentinel (social condition) or a toy mouse (object condition) and allowed to interact for 10 min before being returned to the home cage. After 1 h, the same test mouse was placed back in the same box, which now included either a novel sentinel mouse (social condition) or a novel toy mouse (object condition), and allowed to interact for 10 min before being returned to the home cage. Test mice were anesthetized with ketamine (100 mg/kg, i.p.) 45 min after the last interaction and perfused for subsequent immunostaining. Ventral hippocampal sections from mice containing RetroBeads were stained for the immediate early gene c-Fos and NeuN. mPFC sections from non-surgical mice were stained for c-Fos. All sections were processed for immunohistochemistry at the same time, and images were taken consecutively using identical imaging settings in a confocal microscope (see below).

#### Imaging and quantification of presynaptic terminals

P20 mice received injections of AAV2-Syn-mCherry (UNC Vector Core, Chapel Hill, NC) either in the vHIP (3.8 A/P, 3.3 M/L, 3.5 D/V from the bregma; 500 nL) or in the right mPFC (1.45 A/P, 0.5 M/L, 1.45 D/V from the bregma; 250 nL). They were perfused 4 weeks later, and 30 μm sections were cut from the mPFC and stained with anti-mCherry antibodies. All images were acquired using a 63X (1.4 NA) oil immersion objective in an LSM-800 Airyscan confocal microscope (Zeiss, Oberkochen, Germany) using identical settings (laser power, pinhole, photomultiplier tube, current, gain, and offset). Axons were semi-manually traced using *NCTracer* in *FIJI* (Longair et al., 2011), and boutons were quantified using *BoutonAnalyzer* (Gala et al., 2017) running in Matlab (MathWorks, Natick, MA).

### Electrophysiology

#### Ex vivo brain slices

Mice were anesthetized with ketamine (100 mg/kg, i.p.) and transcardially perfused with an ice-cold modified “cutting” artificial cerebrospinal fluid (aCSF) containing 87 mM NaCl, 2.5 mM KCl, 0.5 mM CaCl_2_, 7 mM MgCl_2_, 1.25 mM NaH_2_PO_4_, 25 mM NaHCO_3_, 25 mM glucose, and 75 mM sucrose, bubbled with 95% O_2_ /5% CO_2_. Coronal sections (300 μm thick) of the mPFC cut at an angle of 10° from the coronal plane were prepared using a vibratome (VT1200S; Leica Biosystems, Wetzlar, Germany), transferred to a submerged recovery chamber filled with “recording” aCSF (125 mM NaCl, 2.5 mM KCl, 2 mM CaCl_2_, 1 mM MgCl_2_, 1.25 mM NaH_2_PO_4_, 25 mM NaHCO_3_, and 25 mM glucose, bubbled with 95% O_2_/5% CO_2_), kept at 30 °C for 30 min, and then allowed to recover for 1 h at room temperature before use.

#### High-speed imaging of voltage-sensitive dye signals

For visualization of vHIP afferent fibers in mPFC slices, P30 mice were injected bilaterally with 0.4 μL of Dextran-Alexa-594 (10,000 MW; Thermo Fisher Scientific, Waltham, MA) at 3.88 A/P, 3.3 M/L, 3.5 D/V from the bregma. After two weeks, mice were sacrificed and individual *ex vivo* slices were stained with 30 μM of the voltage-sensitive fluorescent dye RH414 (Anaspec, Fremont, CA) in aCSF for 1 h at room temperature, and transferred to an immersion chamber continuously perfused (2 mL/min) with “recording” aCSF saturated with 95%O_2_/5% CO_2_ and kept at 30°C. A theta glass electrode filled with “recording” aCSF was positioned within the fluorescently labelled vHIP fiber bundle, and another theta glass electrode was placed in line with the cortical column across from the vHIP fiber electrode. In some experiments, an extracellular electrode filled with “recording” aCSF (1-3 MΩ) was positioned in layer 2/3 to record fEPSPs with an Axoclamp amplifier (Molecular Devices, San Jose, CA) in current-clamp mode; signals were sampled at 2 kHz, amplified 10 times with a pre-amplifier (Model 210; Brownlee Precision, now NeuroPhase, Palo Alto, CA), digitized at 10 kHz (ITC-18; InstruTech, Longmont, CO), and acquired with custom-written software (TI WorkBench) (Inoue, 2018) in a MacMini computer (Apple, Cupertino, CA). RH414 was excited at 530±50 nm with a phosphor-pumped LED (Heliophor; 89 North, Williston, VT), and its filtered fluorescence (535±50 nm band-pass, 580 nm beam-splitter, 594 nm long-pass; Semrock, Rochester, NY) was imaged in an inverted microscope (IX71; Olympus, Tokyo, Japan) through a 10X 0.5 NA objective (Fluar; Zeiss, Oberkochen, Germany) and acquired with a scientific CMOS camera running at 2,500 frames-per-second at full 128×128 pixel resolution (NeuroCMOS-SM128; RedShirt Imaging, Decatur, GA), controlled by NeuroPlex software (RedShirt Imaging, Decatur, GA) in a Windows computer (Dell, Round Rock, TX); the electrophysiology computer was synchronized with the imaging computer by TTL pulses. The input-output relationship of the amplitude of VSD signals was obtained by delivering 12 different stimulus intensities with 30 μA increments. LTP of VSD signals and fEPSPs was induced by high-frequency stimulation of vHIP afferent fibers, which consisted of trains of pulses at 300 Hz for 0.5 s repeated 10 times with 3 min intervals (Huang et al. 2004). Recordings of LTP in slices from symptomatic mice were performed in the presence of 5 μM picrotoxin, whereas those in slices from presymptomatic mice were performed in the presence of 1 μM picrotoxin. VSD signals were analyzed using custom written Matlab codes, as following: data for VSD responses in layer 2/3 were determined in a semi-automated manner with boundaries based on the percent of cortical thickness; the boundary between layers 5 and 2/3 was 20% of the cortical thickness from the white matter, and the boundary between layers 2/3 and 1 was 10% of the cortical thickness from the pia surface. Spatio-temporal spread was calculated as area under the curve (AOC) created by spread in the cortical area (% of total) and time (ms). For the correlation between VSD responses and behavioral performance, only one slice per mouse was used (i.e., the slice with the median response).

#### Intracellular whole-cell recordings

For optogenetic activation of vHIP and contralateral mPFC afferents, a total of 0.5 μL of AAV2-Syn-Chrimson-tdTomato (UNC Vector Core, Chapel Hill, NC) was injected into the left vHIP (250 nL at 3.5 A/P, 3.3 M/L, 3.5 D/V from the bregma and 250 nL at 3.1 A/P, 3.3 M/L, 3.0 D/V from the bregma) and 250 nL of AAV2-Syn-Chronos-GFP (UNC Vector Core, Chapel Hill, NC) into the right mPFC (1.45 A/P, 0.5 M/L, 1.45 D/V from the bregma). After allowing four weeks for opsin expression, mPFC *ex vivo* slices were transferred to an immersion chamber continuously perfused (2 mL/min) with “recording” aCSF (120 mM NaCl, 2.5 mM KCl, 2 mM CaCl_2_, 1 mM MgCl_2_, 25 mM NaHCO_3_, 1.4 mM NaH_2_PO_4_, 21 mM glucose, 0.4 mM Na-ascorbate, and 2 mM Na-pyruvate bubbled with 95% O_2_ / 5% CO_2_) and kept at 30 °C. Addition of 1 μM TTX and 100 mM 4-AP, and increasing Ca2+ to 4 mM ensured recordings of monosynaptic responses (Petreanu et al., 2009). Whole-cell recording electrodes were pulled from borosilicate glass (World Precision Instruments, Sarasota, FL) and filled with 120 mM Cs-gluconate, 17.5 mM CsCl, 10 mM Na-HEPES, 4 mM Mg-ATP, 10 mM NA_2_-creatine phosphate, and 0.2 mM Na-EGTA; this yielded a resistance of 4 ± 0.5 MΩ in aCSF. Whole-cell currents were recorded in voltage-clamp mode with an Axopatch-200B amplifier (Molecular Devices, San Jose, CA), filtered at 2 kHz, digitized at 10 kHz (ITC-18; InstruTech, Longmont, CO), and acquired with TI WorkBench in a G5 PowerMac computer (Apple, Cupertino, CA). Opsins were excited with monochromatic light from a Laser-LED illumination system (Lumen Dynamics, now Excelitas Technologies, Waltham, MA) attached to the epifluorescence port of a DM-LFS fixed-stage upright microscope (Leica Biosystems, Wetzlar, Germany), and focused onto the slice through a Zeiss 63X (1.0 NA) water immersion lens (Zeiss, Oberkochen, Germany), which was also used for visualized whole-cell recordings under infrared-differential interference contrast. TI WorkBench controlled and synchronized the Laser-LED (Lumen Dynamics, now Excelitas Technologies, Waltham, MA) stimulation with electrophysiological recordings and fluorescence imaging with an electron multiplying charge-coupled device (QuantEM 512SC; Photometrics, Tucson, AZ). Chronos was excited with 430 nm light, and Chrimson was excited with 630 nm light. In control experiments, 630 nm light evoked inward currents in layer 5 pyramidal neurons in slices expressing only Chrimson in vHIP axons, but not those expressing Chronos in c-mPFC axons (Figure S9 a-b). Conversely, 430 nm light of an intensity that evoked inward currents in slices expressing only Chronos in c-mPFC axons, did not evoke any responses in slices expressing only Chrimson in vHIP axons (Figure S9b). Neurons were classified as pyramidal or putative interneurons based on their morphology, input resistance, and whole-cell capacitance estimated from the exponential decay of the current response to a test voltage step (Fig S9 c-d). Neurons whose input resistance changed more than 20% during the recordings or had unclamped spikes (i.e., action currents) were not included in the dataset.

### Behavioral assays

#### Three-chamber social assay

Mice were acclimated to being handled during 3 min each for 3 days prior to the testing day at the same time as testing. All handling and testing were done in the dark phase of the 12 h light/ 12 h dark cycle, with the experimenter wearing a red headlamp and infrared illumination for digital videography. Mice were placed in the center chamber of a three-chambered box that contained empty inverted pencil cups in the two side chambers, and were allowed to freely explore. After 5 min of acclimation, mice were shepherded back to the center chamber, and blocks were put in place over the side openings. A novel mouse was put under one of the pencil cups on one of the two sides, with the side being interleaved between trials. The blocks were lifted and the test mouse allowed to freely explore the chambers for 10 min. After this time, the test mouse was again shepherded into the middle compartment and blocked there. A second novel mouse was placed under the previously empty pencil cup. The side blocks were removed and the test mouse was allowed to freely explore for another 10 min. After testing each mouse, the apparatus was thoroughly cleaned with isopropanol. Test mice spending more than 75% of the acclimation time in one compartment were removed from the study.

#### Unrestricted Social Assay

One week prior to testing, the back of sentinel mice was dyed with blond hair dye (Born Blonde Maxi, Clairol) with differing patterns for tracking by computer vision. Mice were acclimated to handling and to the testing box containing clean bedding for 3 days prior to the testing day. All acclimation and testing (3 min handling, 10 min in the testing box) were done in the dark phase with experimenter wearing a red headlamp, and video acquisition using infrared illumination. At the beginning of the testing day, sentinel mice from different cages were placed in a neutral cage to acclimate to one another. The test mice were placed in the testing box and were videotaped for 10 min. Then, sentinel mice (one cage-mate of the test mouse and an unknown from a different cage) were placed in the testing box with the test mouse, and allowed to freely interact while being videotaped for 10 min. After this time, the sentinel mice were placed back in the neutral cage and the test mouse returned to the home cage. The test box was cleaned and filled with new bedding between each test mouse. Each sentinel mice interacted with a maximum of 5 test mice, and were discarded if they fought with other sentinels or were excessively grooming. After all mice had been tested, sentinel mice were individually videotaped for 10 min for computer training. Individual and test videos were fed to the *Motr* program (Ohayon et al., 2013) to create tracks that were sent to *JAABA* (Kabra et al., 2013; Robie et al., 2017) for unbiased computer identification of behaviors. *JAABA* classifiers were trained on pilot data sets. Behavioral scores for social memory assessment were taken from the first 4 min of the trial, and separated based on the mouse target (cage-mate or novel). Behavioral scores for social memory and other behaviors were taken from the entire video and social behavior times pooled between novel and cage-mate mouse. Memory discrepancy scores were calculated as the time spent following the novel mouse divided by time spent following the cage-mate mouse.

#### Novel object recognition

Mice were acclimated to handling and to the testing box containing clean bedding and two identical objects for 7 days prior to the testing day (handled for 3 min, placed in the testing box for 10 min). On the eighth day, mice were returned to the testing chamber where one of the objects had been replaced with a novel object. Time spent interacting with each object was scored using *JAABA*, which was taught to identify active sniffing behavior, and allow quantification of the sniffing time within a user-defined region of interest around each object. Memory scores were calculated for the first 4 min of the test day.

### Chemogenetic manipulation with DREADDs and CNO

#### Chronic treatment

P20 WT and *Mecp2* KO mice were injected with CAV-2-Cre (Biocampus, Institute of Molecular Genetics, Montpellier, France) into the mPFC; 250 nL at 1.45 AP, 0.5 ML, and 1.45 DV, and with either AAV8-hSyn-DIO-mCherry, AAV8-hSyn-DIO-hM4Di(Gi)-mCherry, or AAV8-hSyn-DIO-hM3Dq(Gq)-mCherry into the vHIP; 500 nL at 3.5 AP, 3.3 ML, and 4.0 DV (all viruses from the UNC Vector core). CNO (Tocris Bioscience) was first dissolved in DMSO (5 mg in 200 μL), and then diluted in 200 mL of water with 5 mM saccharine (Sigma Aldrich) (Carvalho Poyraz et al., 2016). Mice were allowed *ad libitum* access to this solution in a standard drinking bottle starting at P34. Mice were tested using the unrestricted social interaction assay at P45. Mice were kept on CNO and sacrificed 3-5 days after behavioral testing for the preparation of *ex vivo* mPFC slices for voltage-sensitive dye imaging.

#### Acute treatment

P20 WT mice were injected with CAV-2-Cre either into the mPFC; 250 nL at 1.45 AP, 0.5 ML, and 1.45 DV; or the NAc; 250 nL at 0.9 AP, 0.9 ML, and 3.8 DV; and AAV8-hSyn-DIO-mCherry or AAV8-hSyn-DIO-hM3Dq(Gq)-mCherry into the vHIP; 500 nL at 3.5 AP, 3.3 ML, and 4.0 DV. P20 *Mecp2* KO mice were injected with CAV-2-Cre into the mPFC; 250 nL at 1.45 AP, 0.5 ML, and 1.45 DV; and AAV8-hSyn-DIO-mCherry or AAV8-hSyn-DIO-hM4Di(Gi)-mCherry into the vHIP; 500 nL at 3.5 AP, 3.3 ML, and 4.0 DV. Mice were acclimated to IP injections for 2 weeks prior to behavioral tests. On testing day, mice received a single IP injection of CNO (3 mg per kg of body weight) 2 h before testing (Smith et al., 2015).

## QUANTIFICATION AND STATISTICAL ANALYSIS

All statistical analyses were performed blinded to the genotype and treatment groups using Prism (GraphPad) and Matlab; Power analyses were performed using G*Power (Erdfelder et al., 2009). Statistical tests used in each experiment are provided in main text, within associated figure legend, or within the statistical table in supplemental information. Significance was defined as p<0.05, with the specific statistical test provided in main text or within associated figure legend in addition to the statistical table in supplemental information. Sample sizes (n) refer to number of cells, number of slices, or number of animals, with the specific convention provided in the main text or within the associated figure legend. Significance conventions are as follows: *: p<0.05; **: p<0.01; ***: p<0.001. Sample sizes are provided in main text, within associated figure legend, or within the statistical table in supplemental information. Behavioral data are presented as mean +/- SEM, all other data are presented as mean +/- SD.

---

**Figure S1, Related to Figure 1.**
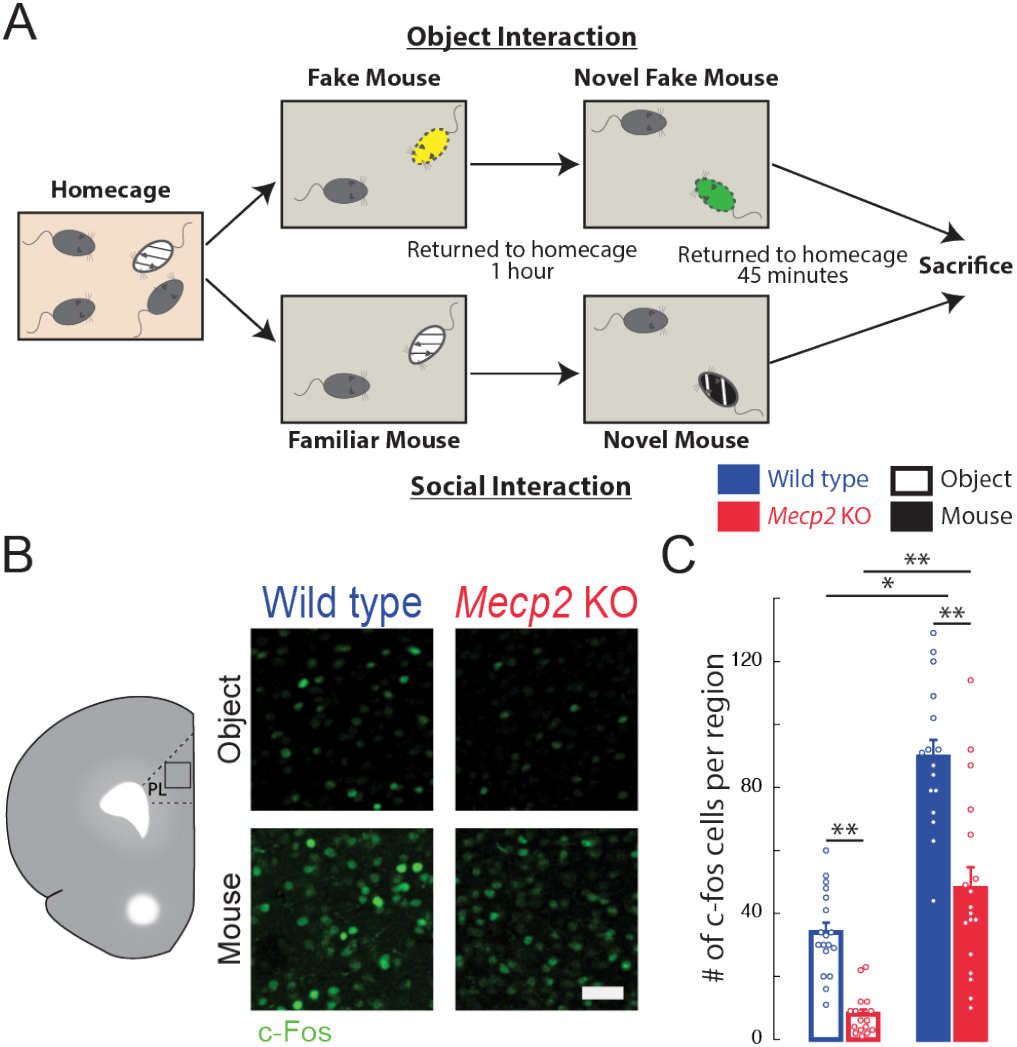
Higher number of c-Fos-positive neurons in the mPFC following social encounters. **(A)** Representative images of c-Fos immunostaining in the mPFC. **(B)** Number of c-Fos positive neurons in the PL region of the mPFC of mice who had interacted with either novel objects or novel mice 45 minutes prior to sacrifice (WT object vs. *Mecp2* KO object, p=0.0073; WT object vs. WT mice, p=0.0008; *Mecp2* KO object vs. *Mecp2* KO mice, p=0.0001; WT mice vs. *Mecp2* KO mice, p=0.0298; Kruskal-Wallis with Dunn’s multiple comparisons; n=9 mPFC sections from three mice in each condition). Scale bar 50 μm. Mean±SEM; * p<0.05, ** p<0.01.

**Figure S2, Related to Figure 3.**
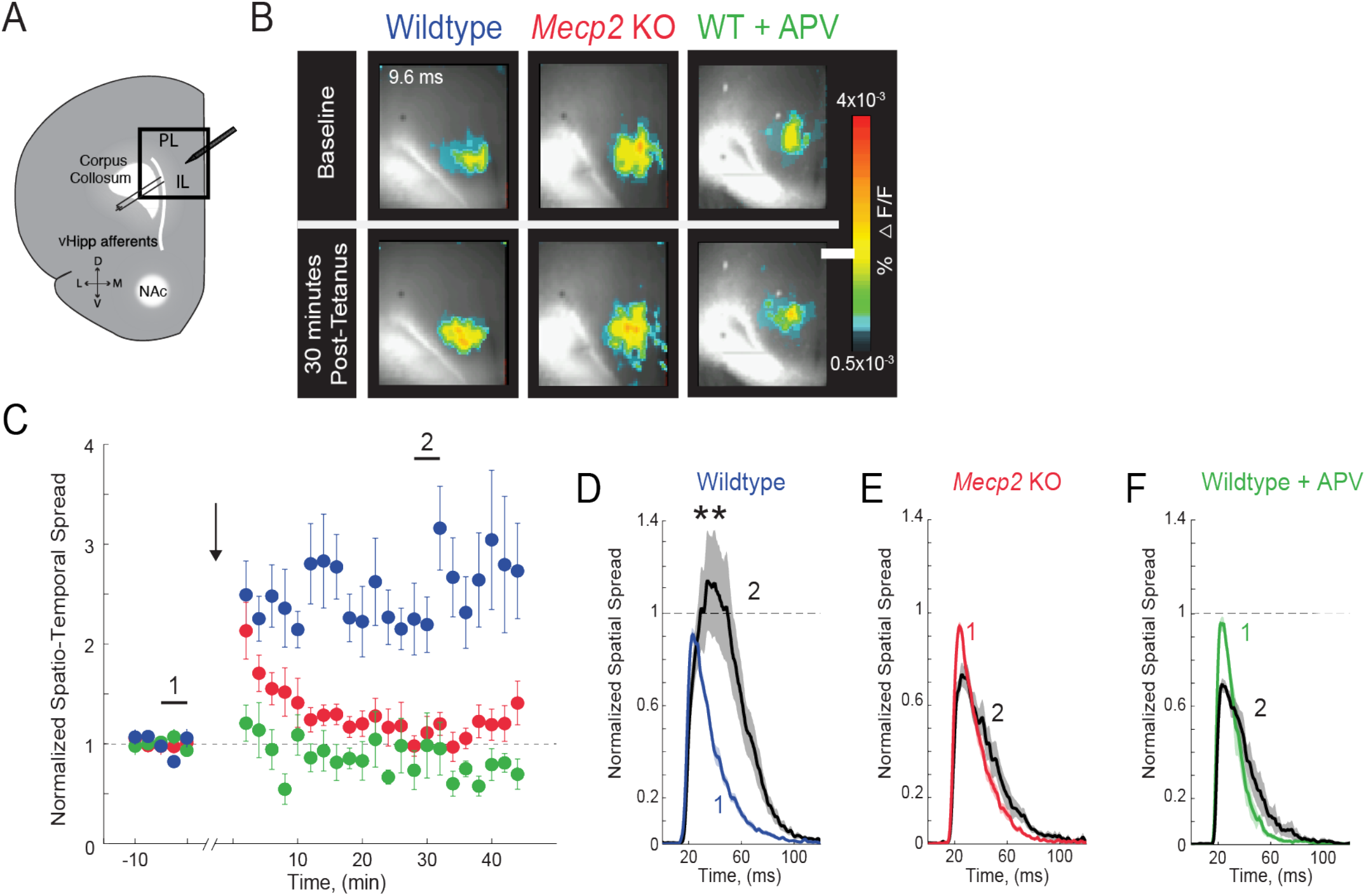
Impaired LTP at vHIP-mPFC synapses in *Mecp2* KO mice. **(A)** Schematic of VSD imaging in an mPFC slice during stimulation of vHIP fibers. **(B)** Representative images of peak VSD responses to a single test stimulation before and after high-frequency stimulation to induce LTP. Scale bar 100 μm. **(C)** Normalized spatiotemporal spread of single test VSD responses before and after LTP induction; high-frequency stimulation given at arrow. **(D-F)** Spread over time of single test VSD responses at baseline (1) and 30 min after LTP induction (2), normalized to baseline spread in WT slices (**D**), *Mecp2* KO slices (**E**), and WT slices in the presence of the NMDA receptor antagonist APV (25 μM) (**F**) (p=0.0013 Student’s paired t-test, n=10 slices from 7 WT mice; p=0.2705 Student’s paired t-test, n=9/5 *Mecp2* KO mice; p=0.9205 Student’s paired t-test, n=4/4 WT mice in APV). Mean ± SEM; * p<0.05, ** p<0.01.

**Figure S3, Related to Figure 4.**
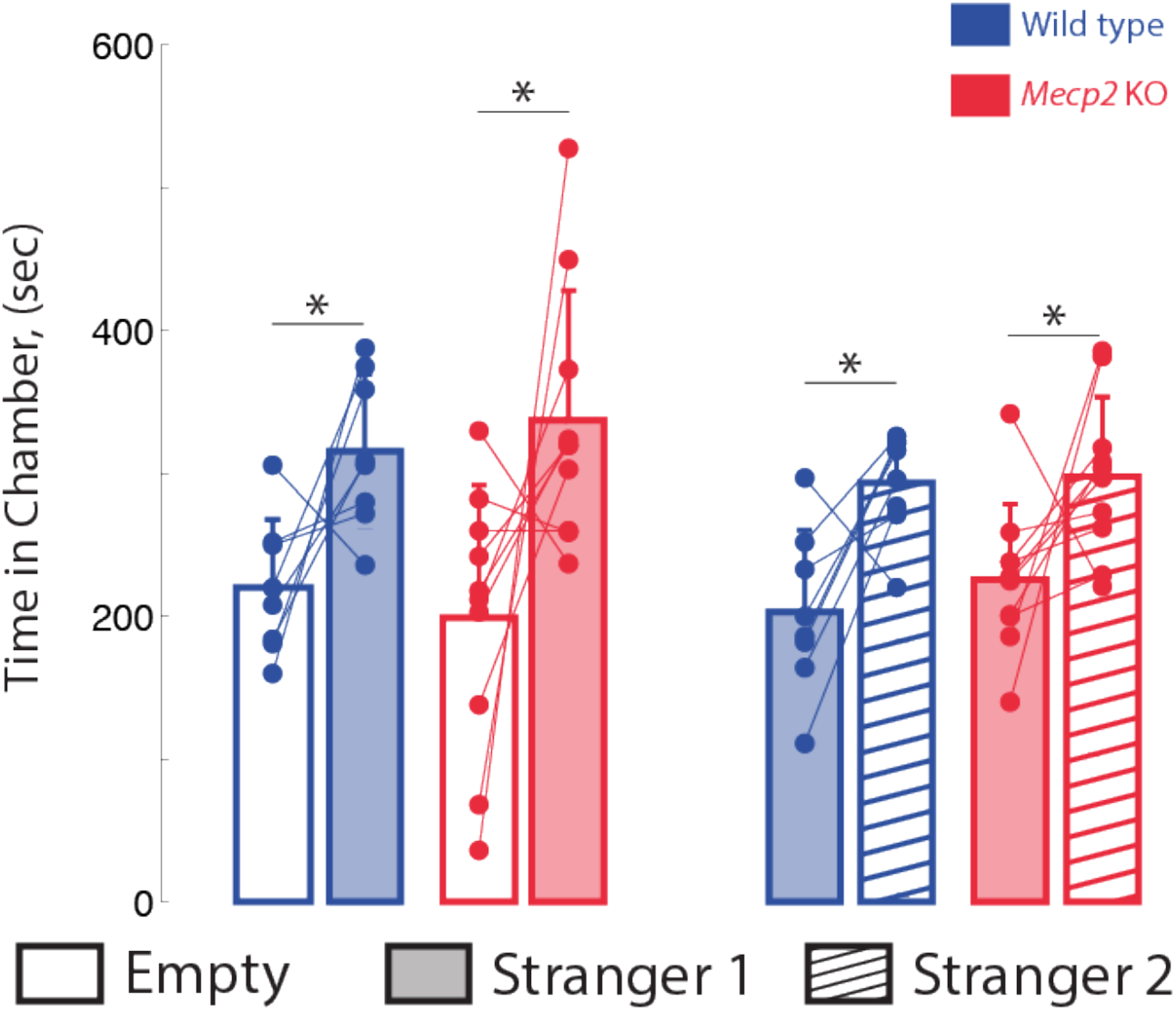
Social memory is not altered in presymptomatic *Mecp2* KO mice. Time spent in chambers with the empty inverted pencil cup, or with a cup containing either stranger 1 or stranger 2 (WT empty vs. stranger 1, p=0.0273; *Mecp2* KO empty vs. stranger 1, p=0.0375; WT stranger 1 vs. stranger 2, p=0.0142; *Mecp2* KO stranger 1 vs. stranger 2, p=0.0484; Student’s paired t-test; n=8 WT mice; n=10 *Mecp2* KO). Mean ± SD; * p<0.05, ** p<0.01.

**Figure S4, Related to Figure 4.**
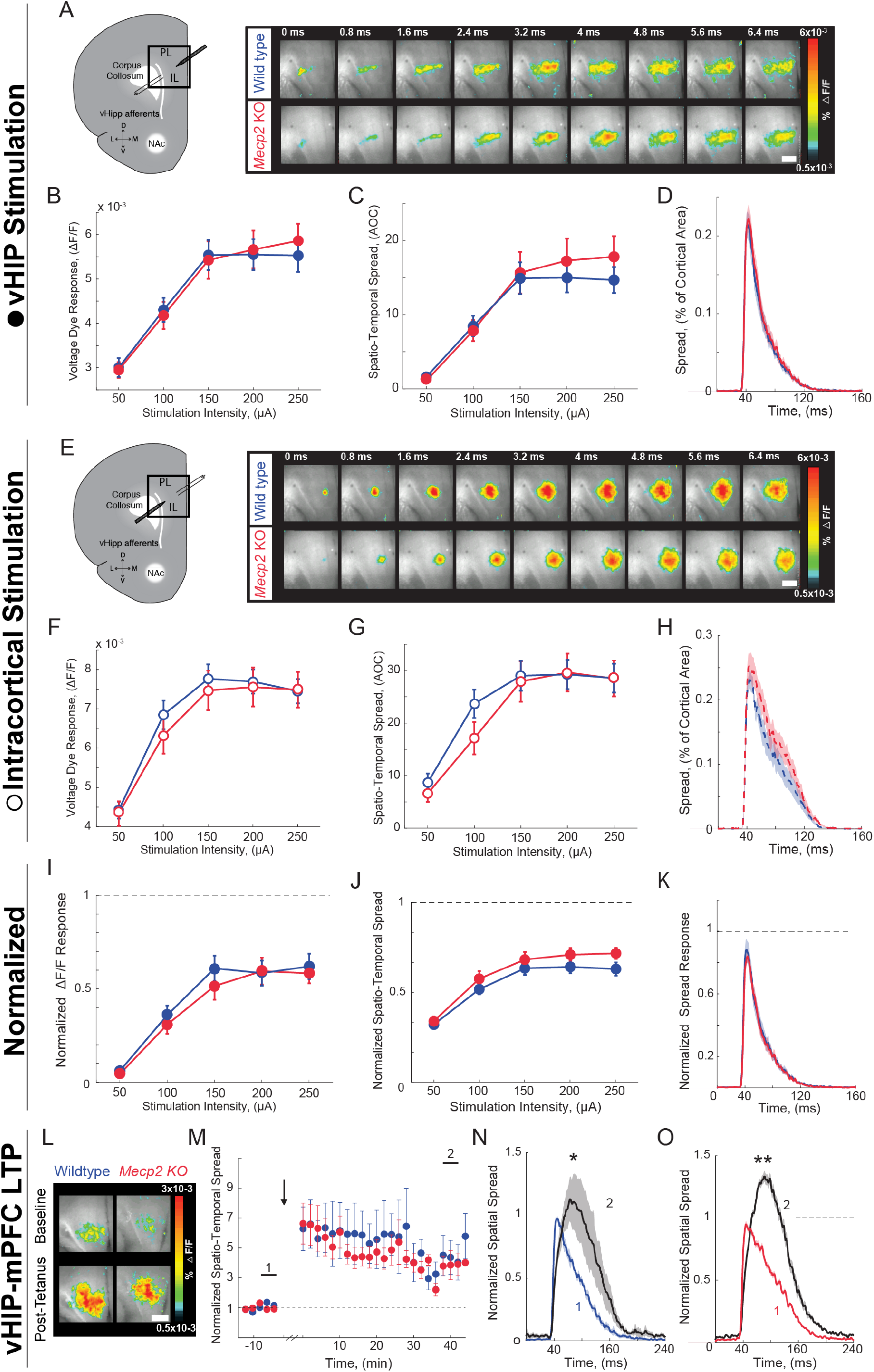
VSD responses to vHIP stimulation in mPFC slices and LTP at vHIP-mPFC synapses are not altered in presymptomatic *Mecp2* KO mice. **(A)** Schematic of VSD imaging in an mPFC slice during stimulation of vHIP fibers and representative VSD responses evoked by a single vHIP fiber stimulation. Scale bar 100 μm. **(B-D)** Input-output relationships of VSD peak responses (**B**, p=0.424, Two-way ANOVA), spatiotemporal spread (**C**, p=0.625, Two-way ANOVA), and spread over time (**D**, p=0.2918 Student’s t-test) (at 150 μA intensity) evoked by a single vHIP fiber stimulation. **(E)** Schematic of VSD imaging in an mPFC slice during stimulation of intracortical fibers and representative VSD responses evoked by a single intracortical stimulation. **(F-H)** Input-output relationships of VSD peak responses (**F**, p=0.4663, Two-way ANOVA), spatiotemporal spread (**G**, p=0.3972, Two-Way ANOVA), and spread over time (**H**, p=0.2582, Student’s t-test) (at 150 μA intensity) evoked by a single intracortical stimulation. **(I-K)** Peak VSD responses (**I**, p=0.2278, Two-way ANOVA), spatiotemporal spread (**J**, p=0.5857, Two-way ANOVA), and spread over time (**K**, p=0.9145, Student’s t-test) evoked by a single vHIP fiber stimulation normalized to those evoked by a single intracortical stimulation. (**A-K** n=14 slices from 5 mice WT; n=18 5 *Mecp2* KO mice). **(L)** Representative images of peak VSD responses to a single test stimulation before and after high-frequency stimulation to induce LTP. **(M)** Normalized spatiotemporal spread of single test VSD responses before and after LTP induction; high-frequency stimulation given at arrow. **(N-O)** Spread over time of single test VSD responses at baseline (1) and 40 min after LTP induction (2), normalized to baseline peak spread in WT slices (**N**, p=0.0159, Wilcoxon paired test) and *Mecp2* KO slices (**o** p=2.165 x10-5, Wilcoxon paired test) (**L-N**, n=5 slices from 5 WT mice; n=9/5 *Mecp2* KO mice). Spatiotemporal spread is the AOC created by spread of the cortical area (% of total) and time (ms). Mean±SEM; * p<0.05, ** p<0.01.

**Figure S5, Related to Figure 4.**
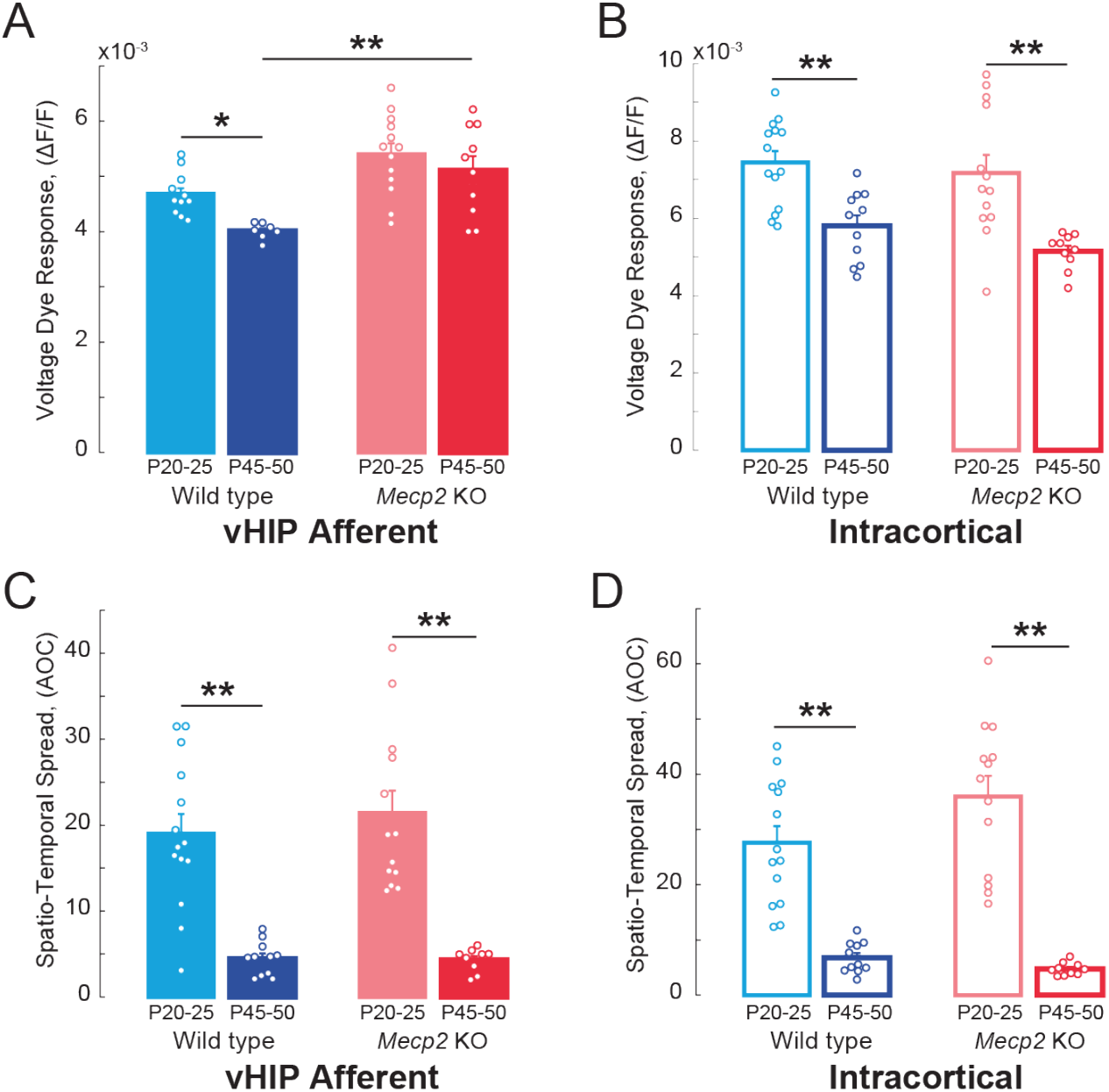
VSD responses to vHIP stimulation in mPFC slices of *Mecp2* KO mice do not develop in a typical manner. **(A-B)** Peak VSD responses evoked by either (**A**) a single vHIP fiber stimulation or (**B**) a single intracortical stimulation in mPFC slices from WT and *Mecp2* KO mice at P20-25 and P45-50. (WT young vs. WT adult vHIP p=0.0440; WT young vs. *Mecp2* KO young vHIP p=0.2378; *Mecp2* KO young vs. *Mecp2* KO adult vHIP p=0.6364; WT adult vs. *Mecp2* KO adult vHIP p=0.0093; WT young vs. WT adult intracortical p<0.001; WT young vs. *Mecp2* KO young intracortical p>0.9999; *Mecp2* KO young vs. *Mecp2* KO adult intracortical p<0.001; WT adult vs. *Mecp2* KO adult intracortical p=0.9999). **(C-D)** Spatio-temporal spread of VSD signals evoked by (**C**) vHIP afferent stimulation or (**D**) intracortical stimulation in mPFC slices from WT and *Mecp2* KO mice at P20-25 and P45-50 (WT young vs. WT adult vHIP p=0.0052; WT young vs. *Mecp2* KO young vHIP, p=0.8426; *Mecp2* KO young vs. *Mecp2* KO adult vHIP, p=0.0032; WT adult vs. *Mecp2* KO adult vHIP, p=0.7319; WT young vs. WT adult intracortical, p<0.001; WT young vs. *Mecp2* KO young intracortical, p=0.9690; *Mecp2* KO young vs. *Mecp2* KO adult intracortical p<0.001; WT adult vs. *Mecp2* KO adult intracortical, p=0.6891) (n=14 slices from 5 P20-25 WT mice; n=18/5 P20-25 *Mecp2* KO, n=11/7 P45-55 WT mice, n=11/5 P45-55 *Mecp2* KO mice; One-way ANOVA with Tukey’s multiple comparisons. Spatiotemporal spread = AOC created by spread of the cortical area (% of total) and time (ms). Mean±SEM; * p<0.05, ** p<0.01.

**Figure S6, Related to Figure 4.**
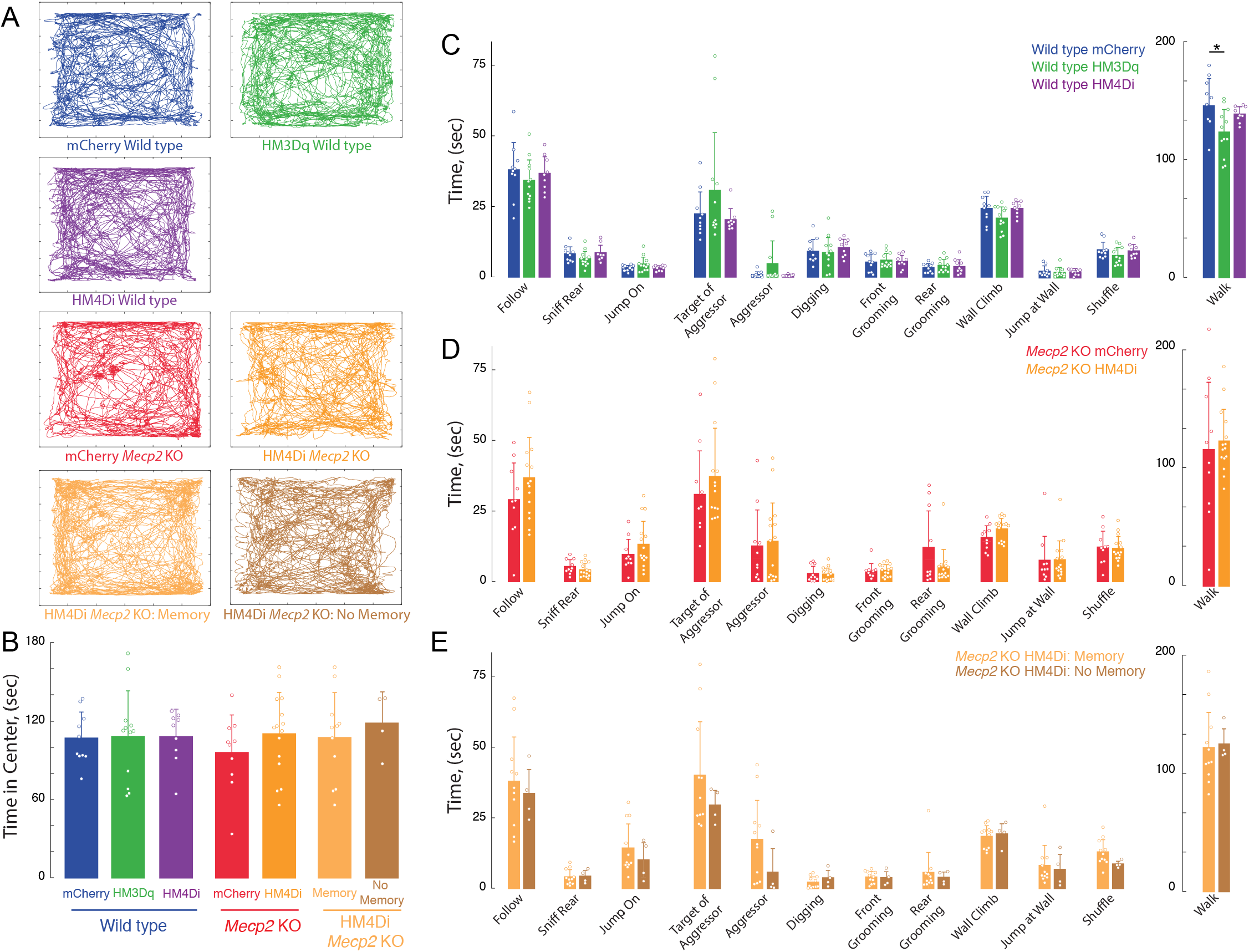
Lack of effects of chronic DREADD stimulation by CNO on anxiety and general (non-social memory) behaviors. **(A)** Representative tracks of individual trajectories of test mice during social encounters in an open field. **(B)** Average time spent in the center of the open field, as a measure of anxiety (mCherry vs. hM3Dq WT p>0.9999; mCherry WT vs. *Mecp2* KO, p=0.9793; hM3Dq WT vs. mCherry *Mecp2* KO, p=0.9566; mCherry vs. hM4D1 WT, p>0.9999; hM4D1 WT vs. mCherry *Mecp2* KO, p=0.9695; hM3Dq WT vs. hM4D1 WT p>0.9999; mCherry vs. hM4D1 (All) *Mecp2* KO, p=0.8910; hM4D1 (All) *Mecp2* KO vs. mCherry WT, p>0.9999; mCherry NAc vs. mCherry mPFC WT, p=0.9492; mCherry vs. hM3Dq NAc WT, p=0.9952; Memory vs. mCherry *Mecp2* KO, p=0.9727; Memory vs. mCherry WT, p>0.9999; No Memory vs. mCherry *Mecp2* KO, p=0.8493; No Memory vs. mCherry WT, p=0.9941; Memory vs. No Memory, p=0.9947; One-way ANOVA with Turkey’s multiple comparisons). **(C-E)** Time spent engaging in social, aggressive, general, and locomotor behaviors during the unrestricted social interaction assay in **(C)** DREADD-expressing WT mice, **(D)** *Mecp2* KO mice (**E**), and hM4D1 *Mecp2* KO mice separated by memory performance (**E**) after chronic CNO treatment (n=9 mCherry WT mice; n=12 hM3Dq WT mice; n=9 hM4D1 WT mice; n=10 mCherry *Mecp2* KO; n=15 hM4D1 *Mecp2* KO mice; Student’s t-test and Kolmogorov-Smirnov test). Mean±SD; * p<0.05, ** p<0.01.

**Figure S7, Related to Figure 5.**
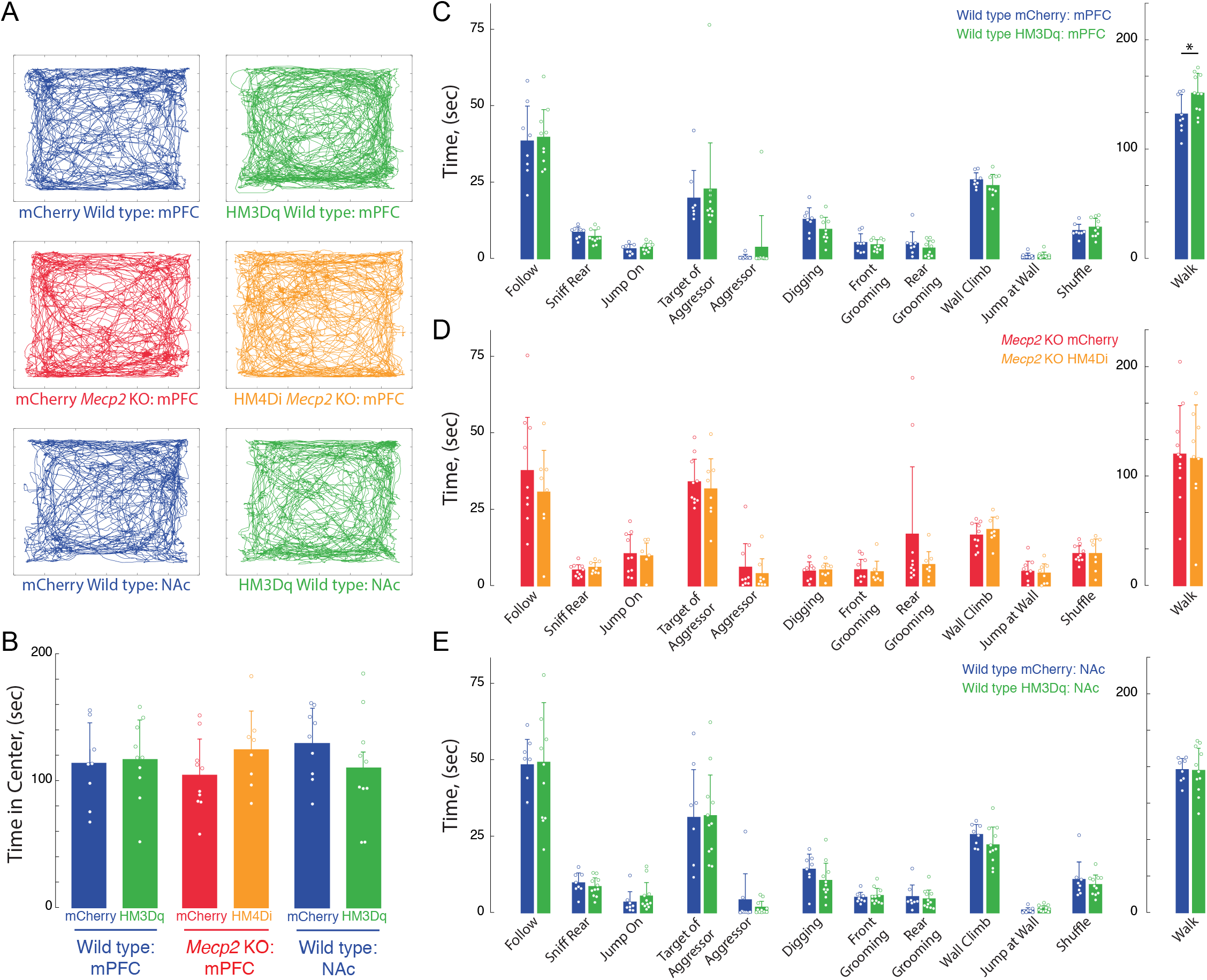
Lack of effects of acute DREADD stimulation by CNO on anxiety and general (non-social memory) behaviors. **(A)** Representative tracks of individual trajectories of test mice during social encounters in an open field. **(B)** Average time spent in the center of the open field, as a measure of anxiety (mCherry mPFC vs. mPFC hM3Dq WT, p>0.9999; mCherry mPFC WT vs. *Mecp2* KO, p=0.9937; hM3Dq WT mPFC vs. mCherry mPFC *Mecp2* KO, p=0.9718; mCherry vs. hM4D1 mPFC *Mecp2* KO, p=0.8502; hM4D1 mPFC *Mecp2* KO vs. mCherry mPFC WT, p=0.9916). **(C-D)** Time spent engaging in social, aggressive, general, and locomotor behaviors during the unrestricted social interaction assay in **(C)** vHIP-mPFC DREADD-expressing WT mice, (**D**) and vHIP-mPFC *Mecp2* KO mice (n=8 vHIP-mPFC mCherry WT mice; n=10 vHIP-mPFC hM3Dq WT mice; n=11 vHIP-mPFC mCherry *Mecp2* KO; n=8 vHIP-mPFC hM4D1 *Mecp2* KO mice). (E) Time spent engaging in social, aggressive, general, and locomotor behaviors during the unrestricted social interaction assay in DREADD-expressing vHIP-NAc WT mice after acute CNO treatment (n=8 vHIP-NAc mCherry WT mice; n=10 vHIP-NAc hM3Dq WT mice; mCherry NAc vs. mCherry mPFC WT, p=0.9492; mCherry vs. hM3Dq NAc WT, p=0.9952; hM3Dq NAc WT vs. mCherry mPFC WT, p=0.9982; hM3Dq NAc WT vs. mCherry mPFC *Mecp2* KO, p=0.9009; One-way ANOVA with Turkey’s multiple comparisons). **(D,E)** Student’s t-test and Kolmogorov-Smirnov test). Mean±SD; * p<0.05, ** p<0.01.

**Figure S8, Related to Figure 6.**
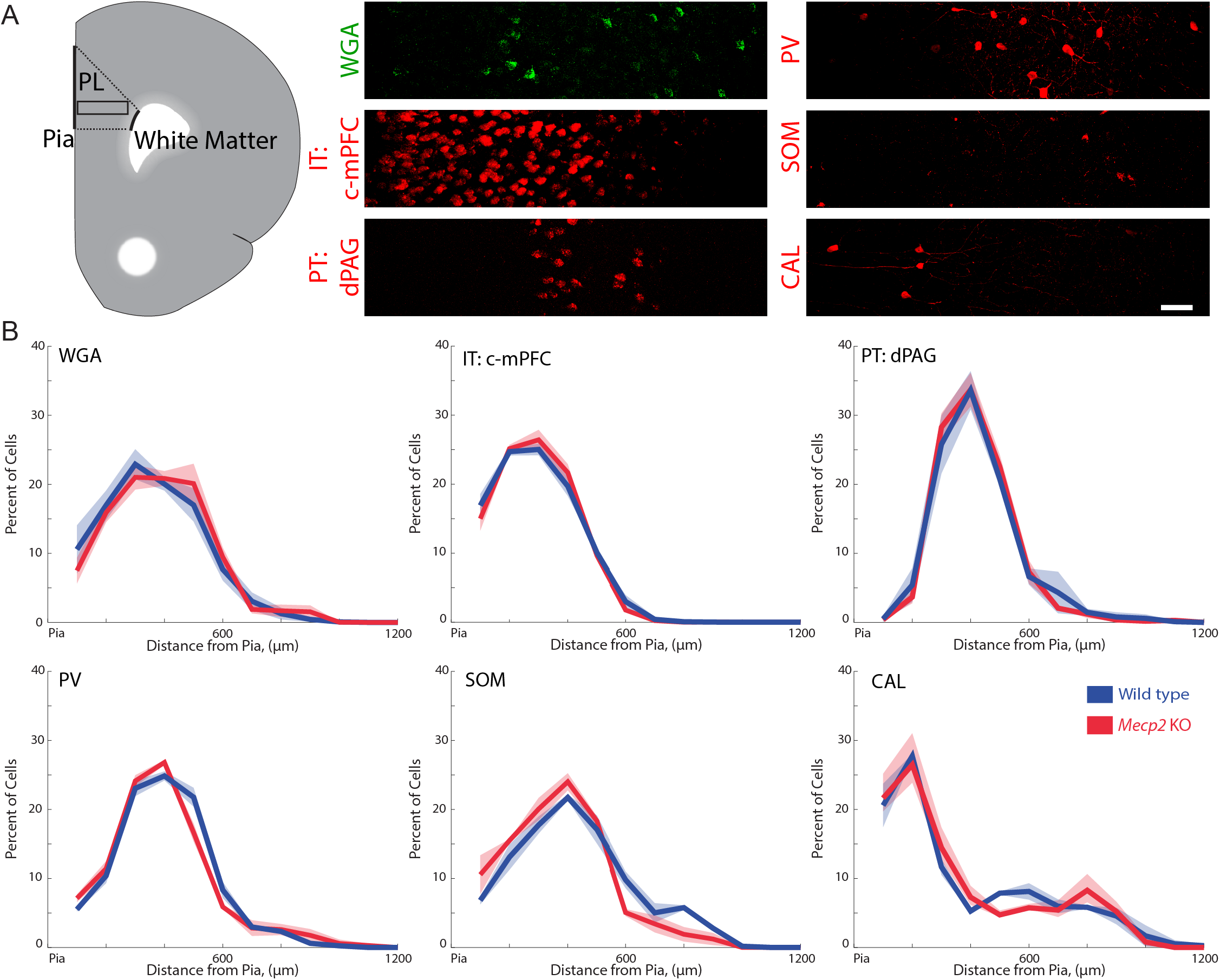
Spatial distribution of different neuronal subtypes in the mPFC that are innervated by vHIP fibers in wild-type and *Mecp2* KO mice. **(A)** Schematic of the PL region of the mPFC (left), and representative images of immunostaining for trans-synaptic WGA; PV, SOM, and CAL interneurons; and retrograde fluorogold labeling of c-mPFC and dPAG pyramidal neurons in the PL-mPFC. Scale bar 50 μm. **(B)** Percent of each neuronal subtype as a function of distance from the pial surface (inter-hemispheric midline) (n=15 sections from 5 mice each for WT and *Mecp2* KO WGA staining, n=9/3 for both WT and *Mecp2* KO mice with c-mPFC, dPAG, PV, SOM, and CAL). Mean±SEM (shaded area).

**Figure S9, Related to Figure 8.**
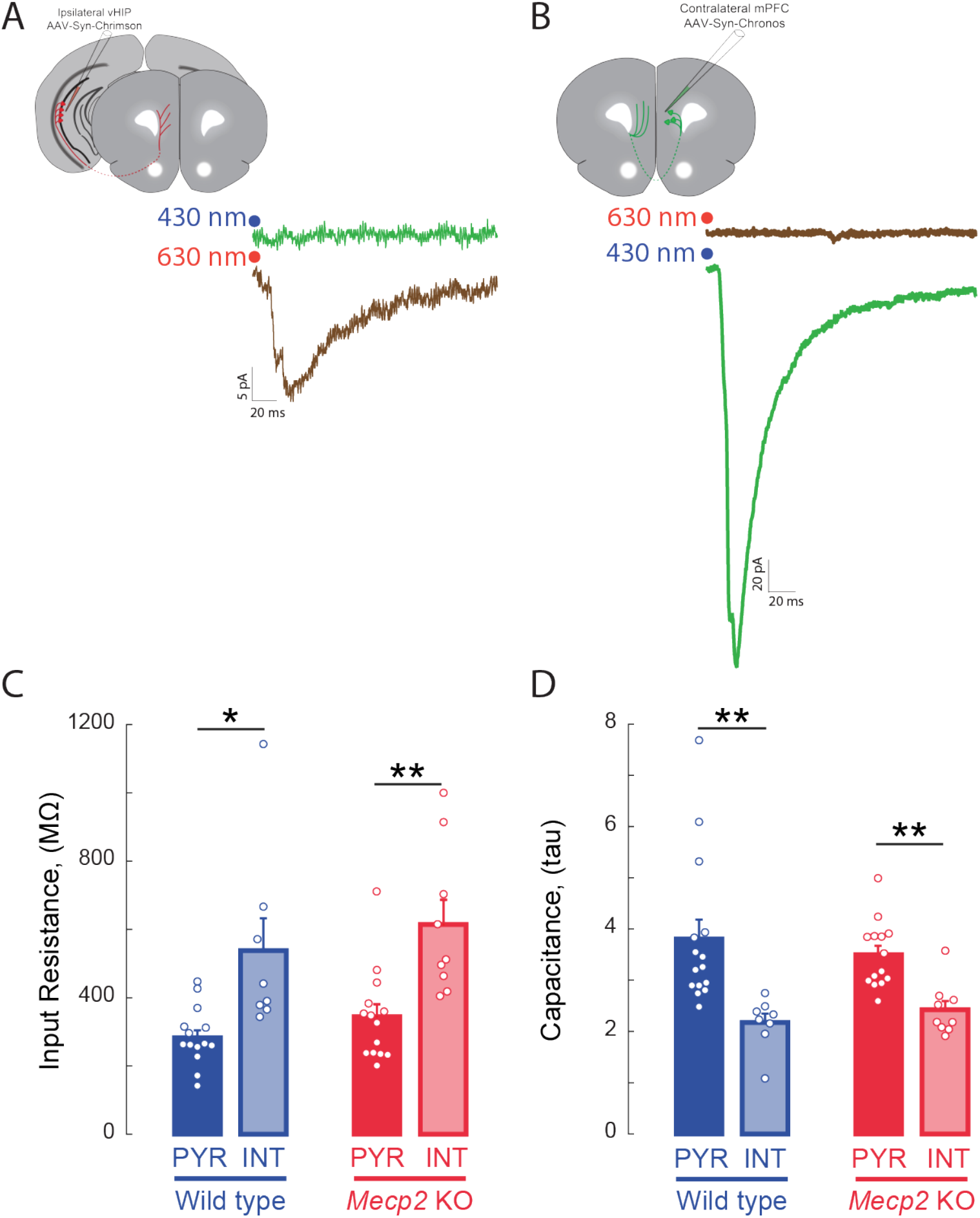
Specific activation of axons expressing opsins with non-overlapping light sensitivity. **(A)** Whole-cell currents evoked by red light (630 nm) in a layer 5 mPFC neuron in a slice from a mouse with Chrimson injection in the vHIP; the same neuron did not respond to blue light (430 nm). **(B)** Whole-cell currents evoked by blue light (430 nm) in a layer 5 mPFC neuron in a slice from a mouse with Chronos injection in the c-mPFC; the same neuron did not respond to red light (630 nm). **(C)** Input resistance and (**D**) whole-cell capacitance of visually identified pyramidal neurons and putative interneurons in layer 5 of mPFC (n=15 pyramidal neurons from 5 slices from 5 WT mice, 8/5/5 WT putative interneurons, 14/5/5 *Mecp2* KO pyramidal neurons, 9/6/6 *Mecp2* KO putative interneurons; p=0.0147 WT pyramidal vs. putative interneurons input resistance, p=0.0083 *Mecp2* KO pyramidal vs. putative interneurons, Kruskal Wallis with Dunn’s Multiple comparisons; p=0.0016 WT pyramidal vs. putative interneurons capacitance, p=0.0035 *Mecp2* KO pyramidal vs. putative interneurons, Kruskal-Wallis with Dunn’s Multiple comparisons). Capacitance estimated from the time constant of the exponential decay of the current response to a test voltage step. Mean±SEM; * p<0.05, ** p<0.01.

